# A ncBAF to cBAF Switch Regulates Human Hematopoietic Progenitor Myeloid Differentiation

**DOI:** 10.64898/2026.10.01.755757

**Authors:** Emmalee R. Adelman, Adedamola A. Adele, Caio Ricardo Gutierrez Silva, Katerina Pilala, Liliana Garcia-Martinez, Oriana Teran Pumar, Felipe Beckedorff, Pradeep Kumar Reddy Cingaram, Rumela Chakrabarti, Defne Bayik, Ramin Shiekhattar, Lluis Morey, Maria E. Figueroa

**Affiliations:** Dept. of Biochemistry, University of Miami Miller School of Medicine, Miami, FL, 33136; Sylvester Comprehensive Cancer Center, University of Miami Miller School of Medicine, Miami, FL, 33136; Dept. of Biochemistry and Molecular Biology, National and Kapodistrian University of Athens, Athens, Greece, 15771; Dept. of Human Genetics, University of Miami Miller School of Medicine, Miami, FL, 33136; Dept. of Molecular and Cellular Pharmacology, University of Miami Miller School of Medicine, Miami, FL 33136; Dept. of Surgical Oncology, University of Miami Miller School of Medicine, Miami, FL 33136

**Keywords:** BRD9, MDSC, bromodomain, hematopoiesis

## Abstract

The mammalian BRG-/BRM-associated factor (BAF) chromatin remodeling complexes are crucial regulators of hematopoietic stem and progenitor cell (HSPC) fate. Expression of the ncBAF subcomponent, BRD9, is altered in both normal aging and myeloid malignancies. We find that in adult human HSPC, BRD9 regulates the differentiation of myeloid progenitors. Upon degradation of BRD9 in these cells, ncBAF is displaced from the chromatin, leading in turn to enhanced recruitment of cBAF to bona fide CEBPA target sites. With the shift from ncBAF to cBAF, HSPC differentiation results in increased generation of myeloid-derived suppressor cells (MDSC) that secrete elevated levels of IL-8. Soluble factors from these myeloid cells, which have lost BRD9, enhance the stemness of breast cancer cells in vitro. In contrast, bromodomain inhibition of BRD9 results in minimal alterations of ncBAF binding and myeloid differentiation, indicating that BRD9 acts as a barrier to human MDSC differentiation in a bromodomain-independent manner.

**KEY POINTS:**

- BRD9 suppresses the development of myeloid-derived suppressor cells in a bromodomain-independent manner
- Loss of BRD9 results in increased cBAF recruitment to CEBPA target sites and enhanced myeloid differentiation

## INTRODUCTION

The BRG-/BRM-associated factor (BAF) (also known as the SWItch/Sucrose Non-Fermentable (SWI/SNF)) chromatin remodelers are modular complexes that regulate nucleosome positioning in an ATP-dependent manner^1^. In mammals, there are three defined complexes, each containing common core proteins such as BRG1, as well as distinct subunits. The canonical BAF (cBAF) complex contains ARID1A/B and DPF2, while the polybromo-associated BAF (PBAF) contains BRD7, ARID2, and PBRM1, and the non-canonical BAF (ncBAF) complex includes GLTSCR1 and bromodomain-containing protein 9 (BRD9)^2^. As there are 29 genes encoding SWI/SNF components and 12-15 proteins per complex, diverse configurations of these regulators are possible^3^. The localization of these complexes plays a critical role in the balance of self-renewal and lineage differentiation of both embryonic and somatic stem cells^4, 5, 6^.

This exquisite control of lineage differentiation through SWI/SNF chromatin remodeling is evident in the hematopoietic system, where loss of individual complex members can result in altered myeloid, erythroid, or lymphoid differentiation^7^. Recently, BRD9 has emerged as an important regulator of murine hematopoietic stem cell (HSC) differentiation^8, 9^. BRD9 is downregulated with normal aging, and loss of BRD9 contributes to the development of aging phenotypes and features of myelodysplastic syndrome (MDS)^8, 10^. In addition to a bromodomain, BRD9 contains a domain of unknown function (DUF), which is essential for interacting with GLTSCR1^11^. Given the role of BRD9 in regulating the multilineage inflammation response^12, 13^ as well as the therapeutic potential of targeting this protein both in solid tumors and hematopoietic malignancies^14, 15, 16, 17, 18, 19^, it is critical to unravel the specific roles of the bromodomain and scaffolding function of BRD9 and its consequences in hematopoiesis.

## METHODS

### Samples

Cryopreserved mobilized peripheral blood CD34+ (HSPC) cells were obtained from the Fred Hutchinson Cancer Research Center, Seattle, Washington. All donors were <35 years old non-smokers. Peripheral blood for T-cell isolation was purchased from Stemcell Technologies. The Institutional Review Board of the University of Miami Miller School of Medicine approved the study.

### ChIP-seq

ChIP-seq was performed with the Ideal TF ChIP Kit (Diagenode, # C01010055)using HSPC that were expanded under hypoxia in IMDM with 20% human serum AB, 100 ng/mL hSCF, 100 ng/mL hTPO 100 ng/mL, 10 ng/mL recombinant hFlt3-ligand, 20 ng/mL hIL-6, and 1% penicillin/streptomycin for 48 hours. Cells from each donor (n=2 per protein) were then split in half and treated with 100 ng/mL dBRD9-A or vehicle control for 72 hours, in hypoxia. Samples treated with vehicle were used for the baseline analysis. Libraries were prepared using the CUTANA Library Prep Kit (Epicypher #14-1001/#14-1002).

## DATA AVAILABILITY

For all data requests, please contact.

## CODE AVAILABILITY

Code used to analyze the datasets in this paper is available upon request.

## RESULTS

### cBAF regulates a distinct set of immune genes in human HSPC

To better understand the specific contribution of BRD9 to epigenomic regulation of human HSPC, we first sought to determine the precise genomic distribution of the three main SWI/SNF complexes in this context. To do this, we performed ChIP-Seq in human CD34+ HSPC for the core component BRG1, and the complex-specific subunits ARID1A and DPF2 for cBAF, ARID2 and BRD7 for PBAF, and BRD9 and GLTSCR1 for ncBAF (n=2 biological replicates) (**Figure 1A and Supplementary Table 1**). Unsupervised analysis of the union of all peaks for all proteins (n=53,173 peaks) revealed that, as expected, binding of each protein correlated most with those of the other proteins within the same complex. Notably, BRG1 binding was most associated with binding of the two PBAF components profiled, BRD7 and ARID2 (**Figure 1B**). Supervised analysis to identify regions bound by cBAF (ARID1A+ or DPF2+), PBAF (ARID2+ or BRD7+), or ncBAF (BRD9+ or GLTSCR1+) complexes in HSPC (**Supplementary Table 2**) showed that the majority of BAF-regulated regions in HSPC are occupied by cBAF (n=48,620), while fewer regions are bound by PBAF (n=19,624) or ncBAF (n=7,229) (**Figure 1C**).

**Figure 1:**
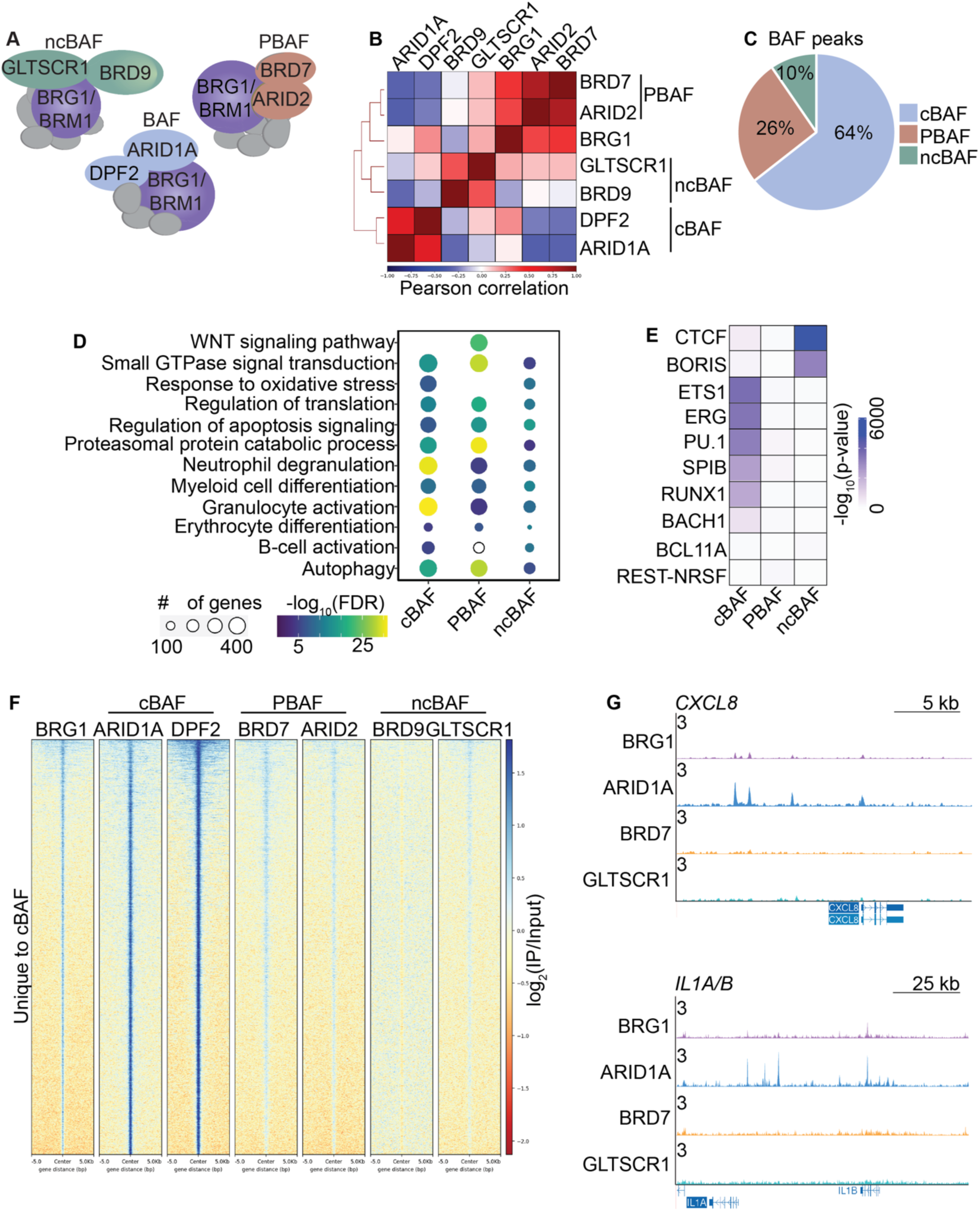
BAF complexes regulate distinct gene sets in human HSPC. **(A)** Schematic representation of the three main SWI/SNF complexes and the subunits studied by ChIP-seq. **(B)** Heatmap of the Pearson correlation for the union of all peaks for each protein that was assayed by ChIP-seq in human HSPC. **(C)** Pie chart of the frequency of each BAF complex. **(D)** Bubble plot of select gene ontology biological processes that are enriched (FDR <0.05) in at least one of the BAF complexes. Circles that are not filled denote non-significant processes. **(E)** Heatmap showing select motifs that are significantly enriched (FDR<0.05) in at least one of the BAF complexes. **(F)** Heatmap of the peaks that are annotated to the genes that are uniquely bound by the cBAF complex compared to PBAF and ncBAF. **(G)** UCSC genome browser tracks at select loci that are uniquely bound by cBAF. Representative tracks from one donor are shown, normalized to counts per million (CPM).

Genomic annotation of SWI/SNF peaks revealed that most PBAF (79%) and ncBAF (54.9%) peaks localize to promoters, whereas cBAF peaks are primarily located in intronic and intergenic regions and exhibit higher enrichment at enhancers (**Supplemental Figure 1A).** ATAC-seq showed that, as a whole, PBAF sites are less accessible compared to cBAF and ncBAF (**Supplemental Figure 1B)**.

Functional annotation showed that cBAF is highly enriched at genes that regulate myeloid activation and differentiation. In contrast, PBAF binds at genes that regulate proteasomal protein processes and WNT and GTPase signaling. While not as enriched as the other complexes, significant binding of ncBAF was also found to be associated with genes in these same pathways (**Figure 1D and Supplemental Figure 1C and Supplementary Table 2)**. Similarly, transcription factor (TF) motif analysis showed significant enrichment of the master myeloid TFs ETS1, PU.1, and RUNX1 at cBAF target sites (**Figure 1E**). PBAF sites showed modest enrichment for these TFs, as well as for REST-NRSF. As expected^8^, the most enriched motif in ncBAF peaks was for the chromosomal architecture-associated protein, CTCF (**Figure 1E**). ChIP-seq of CTCF in HSPC confirmed that 98% of ncBAF and 95% of PBAF peaks overlap with CTCF sites. In contrast, only 49% of cBAF peaks co-localize with CTCF (**Supplemental Figure 1D and Supplementary Table 1**).

Given that many genomic regions have the potential to be bound by various modules (**Supplemental Figure 1E-F**), we next sought to determine whether any genes in human HSPC are solely regulated by a single complex. We identified 3,681 genes that are only regulated by the cBAF complex (**Figure 1F**). These genes include the TFs *CEBPB*, *ETS1,* and *GATA1*, the cytokines *IL1A/B*, *IL3*, *IL5,* and chemokine *CXCL8* (IL-8), and immune signaling genes such as *IFNA1*, *TLR1*, *TLR4*, *and* the alarmins *S100A8/9* (**Figure 1G**). In contrast, we did not identify any genes uniquely regulated by ncBAF or PBAF.

### Loss of BRD9 enhances targeting of cBAF to myeloid genes

To determine how loss of BRD9 impacts BAF complex regulation of HSPC function, we utilized a PROTAC of BRD9, dBRD9-A, that leads to rapid degradation of BRD9, without affecting levels of its homolog BRD7 (**Supplemental Figure 2A-B**). After treating HSPC with dBRD9-A or vehicle for 72 hours, we repeated the ChIP-seq for SWI/SNF components. We identified regions with significant (FDR <0.05) changes in binding of the core (BRG1), cBAF, PBAF, or ncBAF complexes with degradation of BRD9 (**Supplementary Table 3**). As expected, we observed a decrease in ncBAF binding, which was due to both significant reductions in GLTSCR1 (n=2,629 peaks) and BRD9 binding (n=2,271 peaks). PBAF binding appeared to be unperturbed by the degradation of BRD9, with no significant changes detected. Finally, we detected 796 BRG1 peaks gained (306 unique and 490 overlapping with other complexes) and 6,367 cBAF peaks changed (99% gained and 1% lost) (**Figure 2A**). Notably, few of the regions with increased core or cBAF binding upon BRD9 loss correspond to regions marked by ncBAF binding at baseline. Indeed, the overlap of baseline cBAF regions with those having increased binding of the complex after loss of BRD9 suggests that there is both enhanced cBAF binding at its normal target sites and de novo recruitment to 2,391 regions (**Figure 2B-C and Supplemental Figure 2B**).

**Figure 2:**
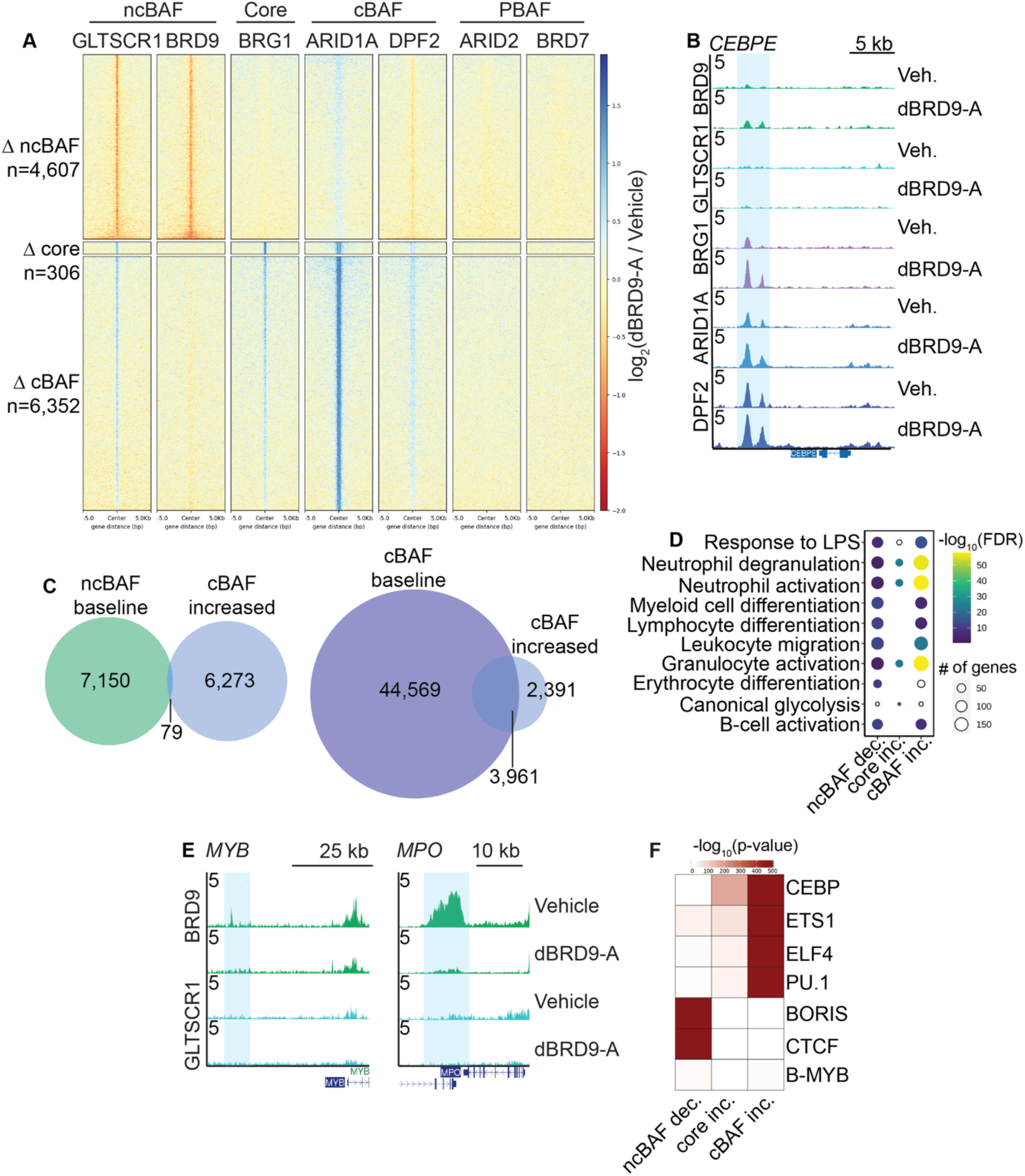
Loss of BRD9 enhances cBAF recruitment to myeloid associated genes. **(A)** Heatmap of genomic regions that have significant (FDR<0.05) changes of core (BRG1), cBAF (ARID1A), or ncBAF (GLTSCR1 or BRD9) binding in HSPC treated with dBRD9-A for 72 hours, compared to vehicle control. Fold-change signal of dBRD9-A over Vehicle is shown for each protein. **(B)** Example of a locus with a significant increase in cBAF binding with loss of BRD9, tracks are from one representative donor and normalized to counts per million. **(C)** Venn diagrams of the overlap of cBAF peaks that are increased with loss of BRD9 with baseline ncBAF peaks (*left*) or baseline cBAF peaks (*right*). **(D)** Bubble plot of select Gene Ontology Biological Processes pathways that are associated with significant (FDR <0.05) changes in BAF binding upon loss of BRD9. Hollow circles denote processes with FDR > 0.05. **(E)** UCSC genome browser tracks of gene loci that are bound by BRD9. Tracks are from one representative donor. **(F)** Heatmap showing the significant enrichment of TF motifs in the regions that have differential BAF complex binding upon loss of BRD9. Heatmap is colored by the -log_10_(p-value) determined by HOMER.

Regions showing increased cBAF or BRG1 were highly enriched for genes associated with granulocyte activation and degranulation as well as the TFs *ETS1*, *KLF4*, *RUNX1*, the chemokines *CXCL1* and *CXCL8*, and alarmins *S100A8/9*. Additionally, there was a significant increase in BRG1 binding at the *CEBPE* locus, a master regulator of granulocyte differentiation. Areas with decreased ncBAF binding upon degradation of BRD9 were associated with myeloid, lymphocyte, and erythroid differentiation and included the TF *MYB* as well as myeloperoxidase (*MPO*) (**Figure 2D-E and Supplementary Table 4**). TF motif analysis of regions with altered BAF complex binding revealed that, as seen at baseline, the areas that have increased core or cBAF binding were enriched for ETS motifs (FDR <0.05). The most significantly enriched motif was for the master family of myeloid TFs, CEBP, which was notably absent at baseline (**Figure 2F**). However, few regions (n=232) had significant changes in CTCF binding with chemical loss of BRD9 (**Supplementary Table 3**).

Next, we asked if changes in chromatin accessibility accompanied BAF binding changes. We performed ATAC-seq on HSPCs treated with vehicle or dBRD9-A for 72 hours (n=5). Analysis of differentially accessible regions (DAR) revealed 608 regions with increased accessibility and 211 with decreased accessibility (**Supplemental Figure 2D and Supplementary Table 5**). Regions with increased accessibility were associated with IL-8 (CXCL8) secretion as well as immune response and were also enriched for CEBP motifs (**Supplemental Figure 2E-F and Supplementary Table 5**). These moderate changes suggest that functional consequences associated with chemical loss of BRD9, are more likely due to changes in binding of chromatin modifiers and transcription factors, rather than broad accessibility alterations.

### CEBPA is overexpressed with loss of BRD9 in a BRD4-dependent manner

Given the significant enrichment of CEBP motifs in regions with increased cBAF binding or accessibility, we performed western blot analysis for members of this family of TFs. We found significant overexpression of CEBPA and CEBPB in HSPC treated with dBRD9-A for 72 hours, while CEBPE was undetectable (**Figure 3A and Supplemental Figure 3A-B**). As CEBPA was the most highly upregulated, we focused on this TF. Protein levels after only 24 hours of treatment with dBRD9-A confirmed that CEBPA overexpression is an early event upon degradation of BRD9 (**Supplemental Figure 3C**).

**Figure 3:**
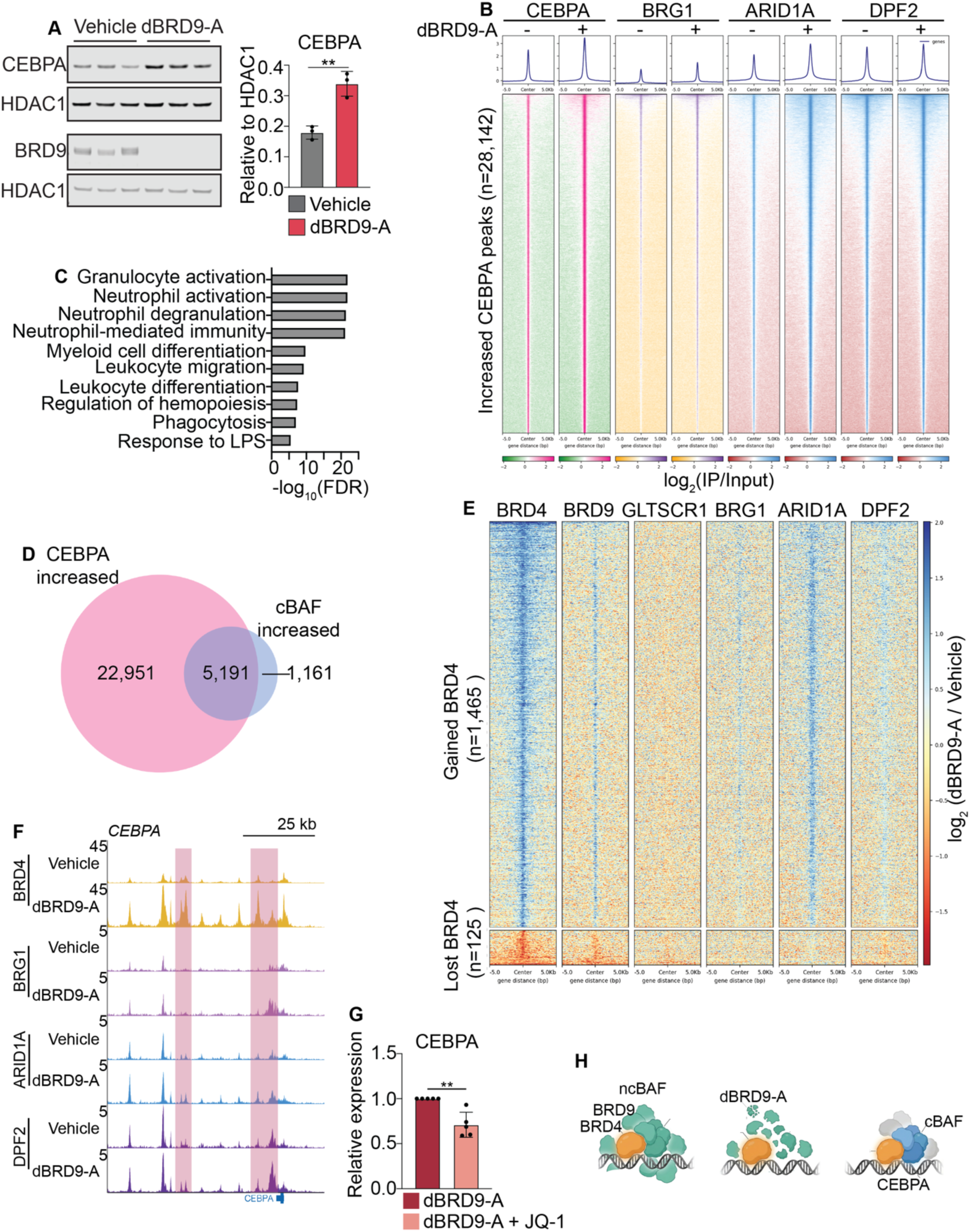
Increased CEBPA expression and localization with cBAF upon loss of BRD9. **(A)** Western blot of CEBPA and BRD9 from HSPC treated with dBRD9-A or vehicle for 72 hours. Loading control is shown for each immunoblot (*left*) and normalization to HDAC1 (*right*), n=3 biological replicates. **(B)**. Heatmap of peaks that have a significant increase in CEBPA with loss of BRD9. The log_2_(Pooled IP/Pooled Input) signal is shown for each protein. **(C)** Bar plot of select significant (FDR < 0.05) GOBP that are enriched in peaks that have increased CEBPA with loss of BRD9. **(D)** Venn diagram of the overlap of peaks with significant (FDR < 0.05) increase in CEBPA or cBAF with loss of BRD9. **(E)** Heatmap of the enrichment of BRD4, BRG1, ARID1A, DPF2, GLTSCR1, and BRD9 binding in HSPC treated with vehicle or dBRD9-A for 72 hours, at regions with significant changes in BRD4 binding upon loss of BRD9. The pooled log_2_(IP/Input) signal is shown for each protein. Error bars represent mean with SD. **(F)** UCSC genome browser tracks of the CEBPA loci. Regions with significant increase of BRD4 binding upon loss of BRD9 are highlighted. Tracks are from one representative donor and normalized to CPM. **(G)** Bar plot of qRT-PCR of CEBPA in HSPC treated with dBRD9-A or dBRD9-A plus JQ-1 for 24 hours. **(H)** Schematic showing the BRD4 dependent regulation of CEBPA by BRD9. Significance calculated using paired-t-test with GraphPad Prism, p>0.05=ns, p ≤ 0.05=*, p≤0.01=**, p≤0.001=***, p≤0.0001=****.

To determine if this increase in CEBPA was accompanied by increased chromatin occupancy, we performed ChIP-seq in HSPC treated with dBRD9-A or vehicle for 72 hours. We observed a significant increase in CEBPA binding at 28,142 regions, with only 55 regions showing decreased binding (**Figure 3B**, **Supplemental Figure 3D and Supplementary Table 3**). As expected, functional annotation revealed that regions with increased CEBPA binding are associated with granulocyte activation and myeloid differentiation (**Figure 3C and Supplementary Table 4**). Notably, 82% of areas with a significant increase in cBAF binding upon loss of BRD9 overlap with sites with increased CEBPA binding (**Figure 3B, D**). Genes in these overlapping regions included the hematopoietic TFs *ETS1*, *RUNX1*, and *ERG*, the alarmins *S100A8/9*, and the chemokines *CXCL1* and *CXCL8* (**Supplemental Figure 3E**).

Given the known association of BRD4 and ncBAF^20^ and that BRD4 can regulate CEBPA expression^21, 22^, we hypothesized that BRD9 regulation of CEBPA may be dependent on BRD4. We first performed Western blot analysis of BRD4 in the chromatin fraction of the human HSPC cell line HUDEP-2 treated with dBRD9-A or vehicle and found a global increase in BRD4 upon degradation of BRD9 (**Supplemental Figure 3F**). ChIP-seq of BRD4 in primary HSPC treated with dBRD9-A or vehicle revealed that, as expected, the majority of baseline ncBAF peaks overlap BRD4 binding sites (**Supplemental Figure 3G**). Yet unexpectedly, upon degradation of BRD9, there is a significant increase in BRD4 binding (n=1,465 peaks), with only a few regions (n=125) losing BRD4 (**Figure 3E**). Notably, the majority (67%) of regions with increased BRD4 binding also had significant increases in cBAF binding (**Figure 3E and Supplemental Figure 3H**) including at the CEBPA locus itself (**Figure 3F**). Combination treatment of HSPC with dBRD9-A and the BRD4 bromodomain inhibitor JQ-1 reduced the effect of dBRD9-A on *CEBPA* expression. Taken together, these results suggest that loss of BRD9 leads to increased targeting of BRD4 and cBAF to the CEBPA locus, which then further recruits cBAF to genes regulating myeloid differentiation (**Figure 3H**).

### BRD9 regulates adult human HSPC stemness in a bromodomain independent manner

To determine whether these changes in cBAF and CEBPA binding were dependent on BRD9’s bromodomain function, we repeated the ChIP-seq for BRD9, GLSTSCR1, BRG1, and ARID1A, as well as CEBPA, in HSPC treated with the vehicle control or the BRD9-specific bromodomain inhibitor, I-BRD9, which has been shown to have similar effects to genetic or chemical loss of BRD9 in other cell types^23, 24, 25^, for 72 hours (**Supplementary Table 6**). No significant differences in the binding of any of the proteins profiled were observed with bromodomain inhibition, either in an unsupervised analysisor when examining the specific regions where we had observed alterations after degradation of BRD9 (**Figure 4A**).

**Figure 4:**
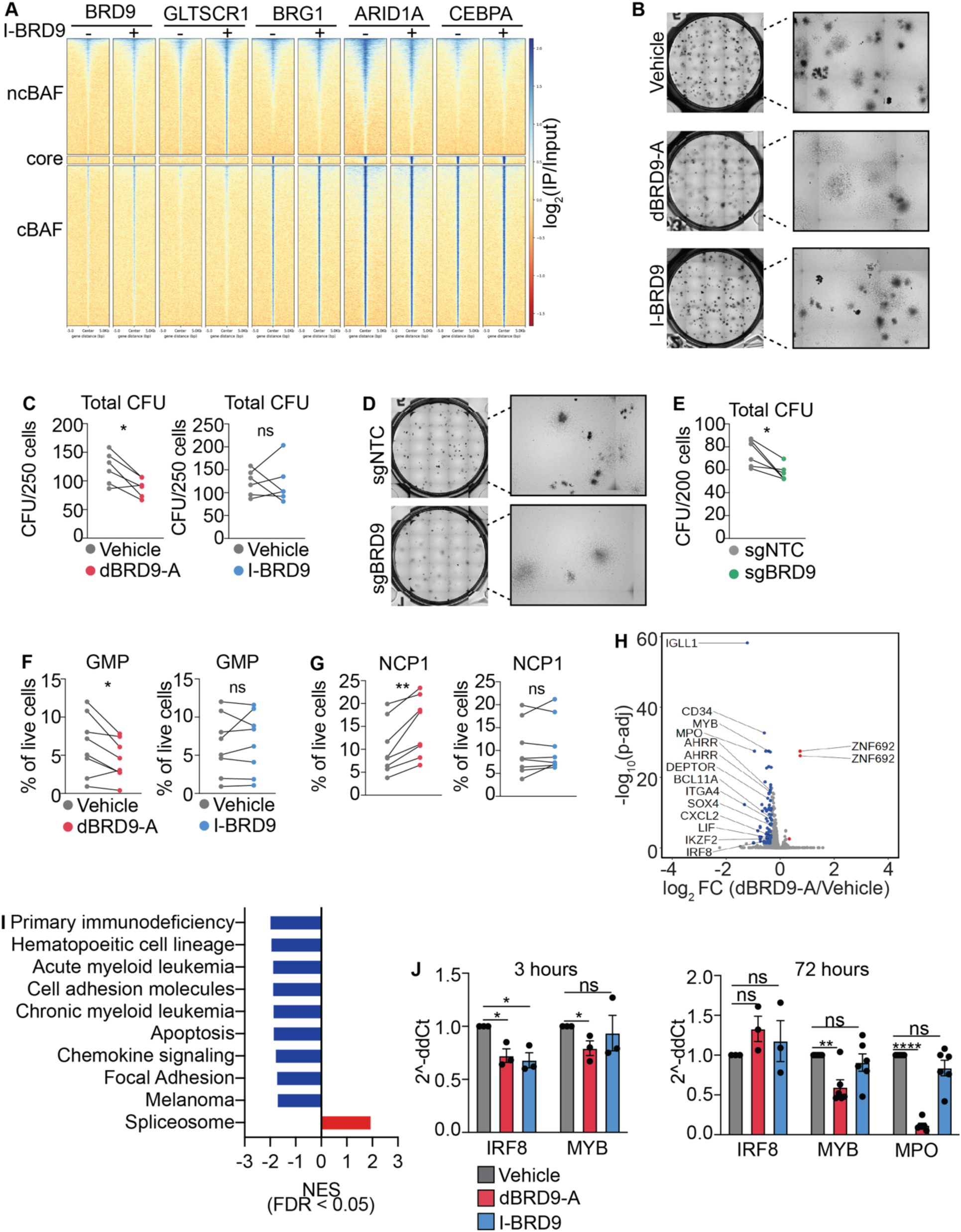
BRD9 regulates HSPC stemness independent of the bromodomain. **(A)** Heatmap of the enrichment of BRG1, ARID1A, GLTSCR1, and BRD9 binding in HSPC treated with vehicle or I-BRD9 for 72 hours, at regions with significant changes in BAF binding upon loss of BRD9. The pooled log_2_(IP/Input) signal is shown for each protein. **(B)** Representative images of CFU and (**C**) plots of the number of total CFU enumerated from plating 250 HSPC treated with dBRD9-A or I-BRD9 for 48 hours (n=6), plotted per donor. **(D)** Representative images of CFU and (**E**) plots of the number of total CFU enumerated from plating 200 CRISPR-Cas9 edited sgNTC or sgBRD9 HSPC 48 hours after electroporation (n=6), plotted per donor. (**F**) Plot of the frequency of GMP and (**G**) NCP1 in HSPC treated with vehicle, dBRD9-A or I-BRD9 (n=8). Error bars represent mean with SD. **(H)** Volcano plot of the PRO-seq transcriptional changes in human HSPC treated with dBRD9-A or vehicle for 3 hr. Significantly (p-adj < 0.05 and absolute foldchange > 1.25) differentially expressed genes are colored in red or blue. **(I)** Bar plot of select significant KEGG pathways that are depleted (blue) or enriched (red) in HSPC treated for three hours with dBRD9-A or vehicle. **(J)** Bar plots of the relative expression calculated by qRT-PCR for select genes after treatment of HSPC with dBRD9-A or I-BRD9 for three hours (*left*) or 72 hours (*right*) (n=3-6). Error bars represent mean with SEM. Significance calculated using paired-t-test with GraphPad Prism, p>0.05=ns, p ≤ 0.05=*, p≤0.01=**, p≤0.001=***, p≤0.0001=****.

Given these expansive changes in cBAF and CEBPA binding, we asked how this would contribute to the altered function of adult human HSPC. We found that degradation of BRD9, but not bromodomain inhibition, either by I-BRD9, or LP99–a dual BRD7 and BRD9–alters colony forming ability. Treatment with dBRD9-A alone results in aberrantly shaped colonies and a reduction in total and erythroid colony-forming units (CFU), with an increase in the percentage of granulocytic-macrophage colonies (CFU-GM) (**Figure 4B-C and Supplemental Figure 4A-B**). To validate the on-target effects of dBRD9-A, we performed CRISPR-Cas9-mediated knockout of BRD9 using guides targeting exon 6 of BRD9 (**Supplementary Figure 4C-D**) and observed that genetic loss of BRD9 also reduced total and erythroid colonies, as well as increased the frequency of CFU-GM (**Figure 4D-E and Supplemental Figure 4E-F**).

We next cultured HSPC under conditions that promote stemness, while treating with dBRD9-A, I-BRD9, LP99, or vehicle for 48 hours and assessed the frequency of granulocyte-macrophage progenitors (GMP, Lineage-, CD66b-, SSC-A low, CD34+, CD45RA+) and neutrophil cell progenitors (NCP) by flow-cytometry^26^ (**Supplemental Figure 5A**). Chemically induced loss of BRD9, but not bromodomain inhibition, resulted in a significant decrease in the frequency of GMP and concurrent increase in NCP1 (Lineage-, CD66b-, SSC-A low, CD34+, CD45RA-, CD115-, CD64dim), which give rise to CD66b+ neutrophils exclusively (**Figure 4F-G and Supplemental 5B-C**). In addition, degradation of BRD9, but not bromodomain inhibition, resulted in decreased NCP2 and NCP4, which are derived from neutrophil monocyte-DC progenitors (**Supplemental Figure 5D**).

We next analyzed nascent transcription using precision run-on sequencing (PRO-seq) on HSPC treated with dBRD9-A or vehicle for three hours. While there are limited early transcription changes at this timepoint (n=82), most of these changes corresponded to downregulated transcripts, including the HSC cell surface antigens *CD34* and *CD38*, as well the hematopoietic TFs *MYB* and *IRF8*, and genes involved in immune signaling, such as *CXCL2*, *LIF,* and *IGLL1*, and *MPO,* a hallmark myeloid enzyme (**Figure 4H**). Evaluation of *MYB* in and *MPO* gene expression in CRISPR-Cas9 edited cells confirmed that they are downregulated with loss of BRD9 (**Supplemental Figure 5E**). Gene set enrichment analysis (GSEA) revealed that dBRD9-A downregulated genes are associated with hematopoietic lineage, myeloid leukemia, and chemokine signaling (**Figure 4I and Supplementary Table 7**). Analysis of gene expression of BRD9 target genes by qRT-PCR revealed that I-BRD9 affects only a subset of these genes and that some of these transcriptional changes persist in dBRD9-A-treated HSPC over time (**Figure 4J**). These results are in line with models of genetic loss of BRD9 in human cord blood CD34+ and murine HSC and suggest that BRD9 regulates neutrophil progenitors in adult human hematopoiesis through regulation of MYB, IRF8, and MPO in a bromodomain-independent manner.

### BRD9 loss and bromodomain inhibition have distinct effects on myeloid differentiation

We next sought to determine if chemical degradation of BRD9 alters normal myeloid maturation. To do this, we treated human HSPC with BRD9 inhibitors for 48 hours, followed by culture in myeloid-promoting media for seven days. HSPC were continuously treated with the different agents, which was well tolerated by the cells (**Figure 5A and Supplemental Figure 6A-B**). Notably, degradation of BRD9, but not bromodomain inhibition, either by I-BRD9 or LP99, markedly altered cell morphology and resulted in increased expression of monocytic and neutrophil-associated cell surface antigens (**Figure 5B and Supplemental Figure 6C-D**). Instead, bromodomain inhibition resulted in only minor changes of cell surface markers (**Figure 5C and Supplemental Figure 6C-D)**. Importantly, differentiation of HSPC in the presence of CFT8634, a second BRD9 degrader, as well as CRISPR-Cas9 knockout of BRD9, showed similar effects on myeloid differentiation, validating these findings (**Figure 5D** and **Supplemental Figure 6E**)^14^.

**Figure 5:**
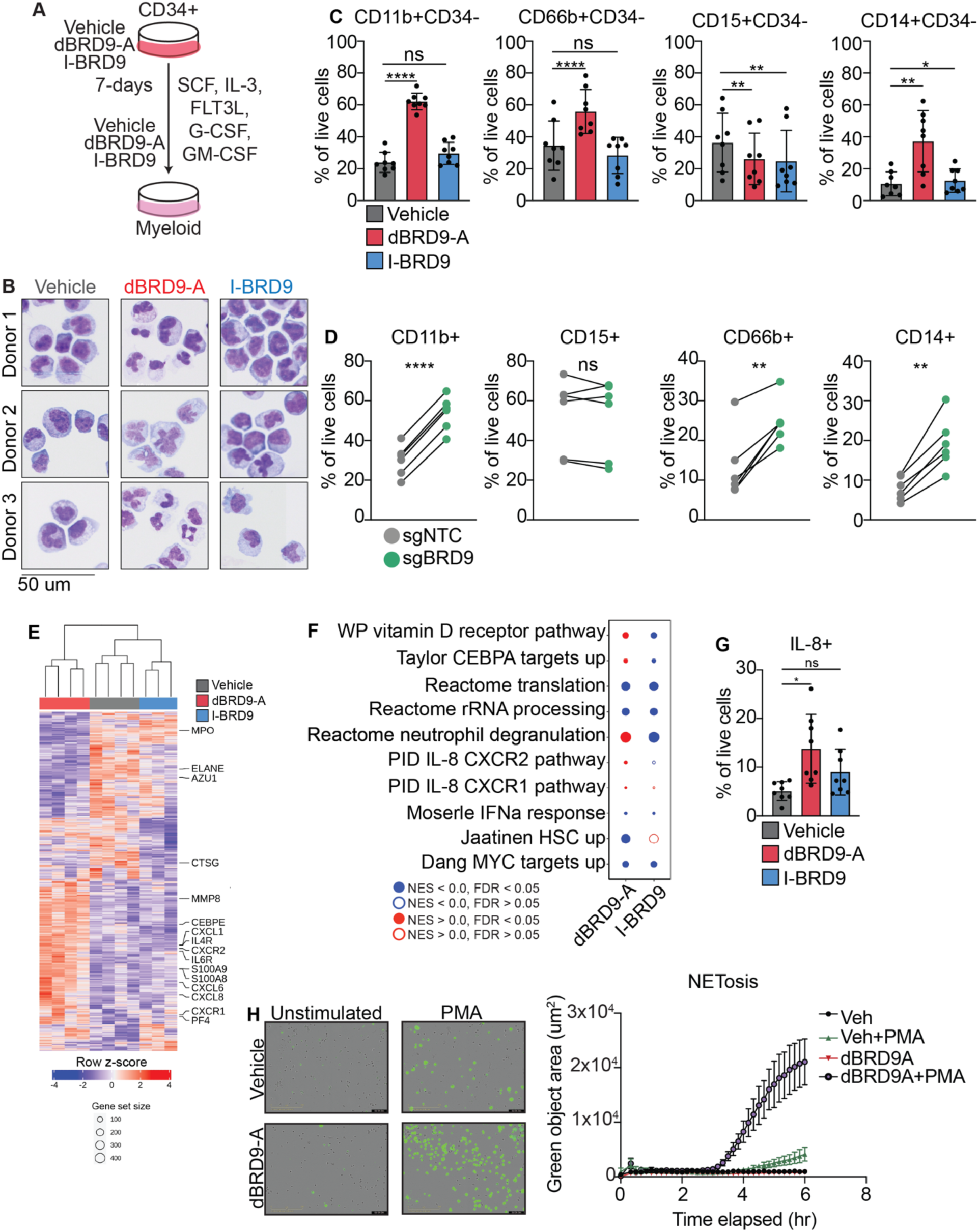
Loss of BRD9 results in highly active IL-8-secreting myeloid cells. **(A)** Schematic of the differentiation of myeloid cells from HSPC with concurrent treatment with BRD9 inhibitors. **(B)** Representative cytospin images from myeloid cells treated for seven days with dBRD9-A, I-BRD9, or vehicle control. **(C)** Summary bar plots of frequencies of cells expressing the given markers, as measured by flow-cytometry on day seven of myeloid differentiation with treatment of BRD9 inhibitors (n=8). Error bars represent mean with SD. **(D)** Summary plots of frequencies of cells expressing the given markers, in sgNTC and sgBRD9 myeloid cells on day seven of myeloid differentiation. **(E)** Heatmap showing all genes that are differentially expressed with BRD9 chemical inhibition compared to vehicle control. Select genes are labeled (n=3-4 donors). **(F)** Bubble plots of select significant (FDR < 0.05) gene set enrichment analysis pathways that are deregulated with dBRD9-A or I-BRD9 treatment in myeloid cells. **(G)** Bar plot of the percentage of live myeloid cells that express intracellular IL-8 after seven days of treatment with dBRD9-A, I-BRD9, or vehicle (n=8). Error bars represent mean with SD. **(H)** Example Incucyte images of dBRD9-A and vehicle myeloid cells stimulated with PMA and stained with cytotox green to measure NETosis (*left*) and quantification of NETosis during the six hours after stimulation (*righ*t) (n=3), error bars are for the mean with SEM. Significance calculated using paired-t-test with GraphPad Prism, p>0.05=ns, p ≤ 0.05=*, p≤0.01=**, p≤0.001=***, p≤0.0001=****.

To investigate the mechanisms underpinning the phenotypic changes in these cells upon inhibition or degradation of BRD9, we performed RNA-seq on myeloid cells that had been continuously treated with the BRD9 inhibitors or vehicle for seven days. Differential gene expression analysis revealed that dBRD9-A and I-BRD9 resulted in distinct transcriptional profiles, with 1,213 differentially expressed genes in dBRD9-A-treated cells, whereas I-BRD9 treatment led to fewer (n=492) transcriptional changes (**Figure 5E**). Direct comparison of differentially expressed genes with either inhibitor revealed only 172 DEG shared between the two, including downregulation of genes associated with primary/azurophilic granules, such as *AZU1*, *ELANE*, and *MPO* (**Figure 5E and Supplemental Figure 6F**). However, upregulation of secondary, tertiary, and secretory granule genes, such as *S100A8/9,* was only observed in dBRD9-A-treated cells. Furthermore, upregulation of immune signaling-related genes *CXCL1*, *CXCL6*, *CXCL8*, and the CXCL8-receptor, *CXCR2*, and the master granulocytic TF, *CEBPE,* was only detected with dBRD9-A (**Figure 5E, Supplemental Figure 6F, and Supplementary Table 8)**.

### Enhanced differentiation of IL-8-secreting cells upon loss of BRD9

To further understand how BRD9 regulates human myeloid cell function, we performed GSEA and found that both I-BRD9 and dBRD9-A treatment resulted in downregulation of genes associated with MYC, a known target of BRD9. However, only dBRD9-A led to upregulation of genes associated with neutrophil degranulation and IL-8 signaling (**Figure 5F and Supplementary Table 8)**. Notably, IL-8 (CXCL8) is a chemoattractant cytokine that is elevated in hematopoietic and solid-tissue cancers^27, 28, 29^. Intracellular flow cytometry analysis confirmed increased expression of IL-8 in dBRD9-A-, but not I-BRD9-treated myeloid cells (**Figure 5G**). Furthermore, an immunoassay confirmed enhanced IL-8 secretion with degradation of BRD9 (**Supplementary Figure 6G**).

As we observed both an increase in IL-8 and its receptors, CXCR1/2, in dBRD9-A myeloid cells, we next asked whether this was associated with an increase in neutrophil-like functions. To investigate this, we stimulated vehicle and dBRD9-A-treated cells with poly (methyl)acrylate (PMA) and measured NETosis^30, 31^. Live imaging revealed that degradation of BRD9 increases NETosis upon PMA stimulation (**Figure 5H**). Together, these results suggest that BRD9 acts as a barrier to myeloid differentiation and loss of BRD9, but not bromodomain inhibition, results in increased differentiation of active IL-8-secreting myeloid cells.

### BRD9 regulates the differentiation of myeloid-derived suppressor cells

Because our work and the research of others suggest that BRD9 inhibits the differentiation of both monocytes and granulocytes, to glean more insight into what specific cell types are affected by loss or inhibition of BRD9, we first used EnrichR and PanglaoDB to identify cell types associated with the genes that are differentially expressed in dBRD9-A or I-BRD9 myeloid cells compared to control. Although both sets of downregulated genes relate to erythroid precursors, B-cells, and T-cells, only BRD9 degradation resulted in myeloid cells upregulating genes associated with neutrophils and myeloid-derived suppressor cells (MDSC) (**Figure 6A**). Flow cytometry analysis of HSPC-derived myeloid cells that had been treated continuously with dBRD9-A, I-BRD9, or vehicle confirmed that degradation of BRD9 leads to a significant increase in the frequency of monocytic MDSC (M-MDSC; CD14+CD15-HLADR-), monocytes (CD14+CD15-HLADR+), and polymorphonuclear MDSC (PMN-MDSC; CD15+CD14-CD11b+CD66b+), while I-BRD9 only affected M-MDSC to a much lesser degree and LP99 did not result in any significant changes (**Figure 6B and Supplemental Figure 7A-B)**. Treatment of myeloid cells with LP99 and CFT8634 also confirmed these findings (**Supplemental Figure 7C-D**) and CRISPR-Cas9-mediated knockout of BRD9 resulted in similar increases of MDSC as the BRD9 PROTACS (**Figure 6C-D**).

**Figure 6:**
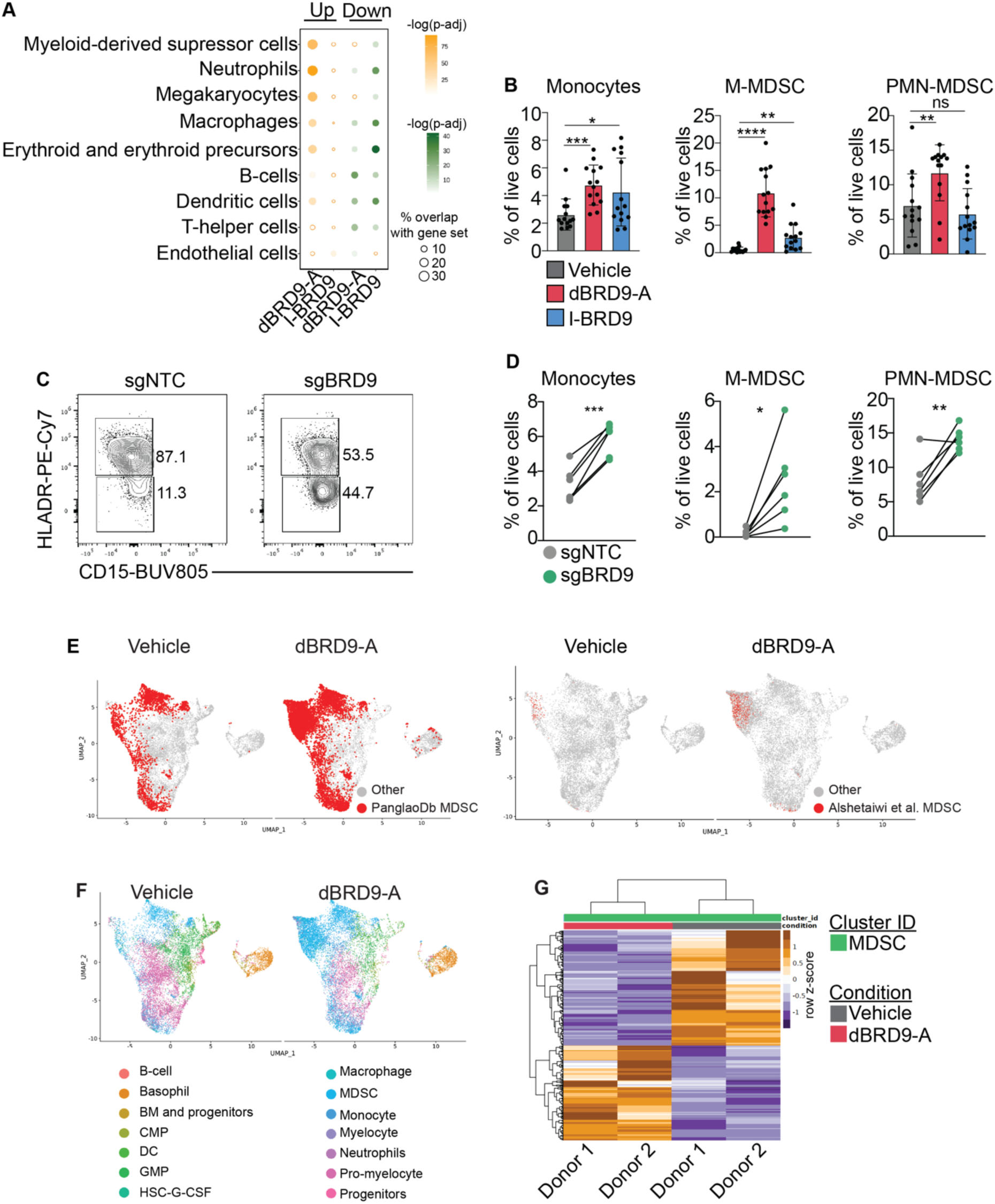
BRD9 regulates the differentiation of myeloid derived suppressor cells. **(A**) Bubble plot of the top cell types (EnrichR) that are associated with genes significantly upregulated (orange) or downregulated (green) with dBRD9-A or I-BRD9 treatment of myeloid cells compared to vehicle. Circles are colored by the -log_10_(p-adjusted); hollow circles denote p-adjusted > 0.05. **(B)** Summary bar graphs of M-MDSC, monocytes, and PMN-MDSC cell frequencies as a percentage of live cells (n=14). Error bars depict mean with SD. Significance calculated using paired-t-test. **(C)** Representative flow-cytometry plots of sgNTC and sgBRD9 monocytes and M-MDSC, on day seven of myeloid differentiation. M-MDSC are gated on CD14+CD15-cells. **(D)** Plots of M-MDSC, monocytes, and PMN-MDSC cell frequencies from sgNTC and sgBRD9 myeloid cells, as a percentage of live cells (n=6). Each point is from one donor. **(E)** UMAP plots of scRNA-seq from myeloid cells treated with dBRD9-A or vehicle. Cells are colored by MDSC status based on the Panglao dB MDSC signature (*lef*t) or published patient MDSC gene signature (*right*), n=2 biological replicates shown per condition. **(F)** UMAP plots of scRNA-seq from myeloid cells treated with dBRD9-A or vehicle. Cells are colored by cellular annotation, n=2 biological replicates shown per condition. **(G)** Heatmap of genes that are significantly differentially expressed in dBRD9-A MDSC compared to vehicle MDSC. Significance calculated using with GraphPad Prism, p>0.05=ns, p ≤ 0.05=*, p≤0.01=**, p≤0.001=***, p≤0.0001=****.

To determine whether these *in vitro*-generated MDSC resemble those found in patients, we performed single-cell RNA-seq (scRNA-seq) in myeloid cells treated with dBRD9-A or vehicle. Using publicly available gene expression signatures of MDSC^32, 33^ and comparing it to our scRNA-seq results, we found that degradation of BRD9 resulted in significant enrichment of cells expressing not only a broad MDSC signature, but also of that of primary breast cancer MDSC, which has been shown to identify human PMN-MDSC (**Figure 6E**). Along with this enrichment of MDSC, we observed an increase in the of macrophage and neutrophil subpopulations and a concurrent decrease in progenitors, confirming our in vitro immunophenotypic analyses (**Figure 6F and Supplemental Figure 7E**). These MDSCs have increased expression of *CXCL8* and the alarmins *S100A8/9/11/12* compared to other cell types (**Supplemental Figure 7F-G**). Pseudobulk differential gene expression analysis revealed 972 genes significantly altered in dBRD9-A MDSC compared to vehicle, including downregulation of *CD24*, a T-cell co-stimulatory factor, and upregulation of *SPARC* (osteonectin), a secreted protein that enhances tumor epithelial-to-mesenchymal transition^34^ (**Figure 6G and Supplementary Table 9**). GSEA analysis of this dBRD9-A MDSC gene expression signature further revealed upregulation of TGFβ, TNFα, and IL-6 signaling pathways, while downregulated genes are associated with MYC targets and DNA repair. These findings demonstrate that these MDSCs are not just quantitatively different, but also qualitatively different from vehicle-treated MDSCs **(Supplementary Table 9)**.

### BRD9 degradation results in the expansion of functional MDSCs

To validate that these *in vitro-*generated M-MDSC maintain known characteristics of this suppressive population, we evaluated levels of reactive oxygen species (ROS) and intracellular TGFb^35^. We found that the majority of CD14+CD15-HLADR-defined M-MDSC contain ROS and TGFb in both vehicle and dBRD9-A M-MDSC, with dBRD9-A treatment resulting in the generation of significantly more ROS+ cells than vehicle, as well as an increase in the frequency of M-MDSC expressing CD84, which is a marker of MDSCs in primary breast cancer^32^ (**Figure 7A**). Similarly, we observed that BRD9 degradation increased the frequency of highly pure LOX1+CD10-PMN-MDSC (**Supplementary Figure 8A-B**).

**Figure 7:**
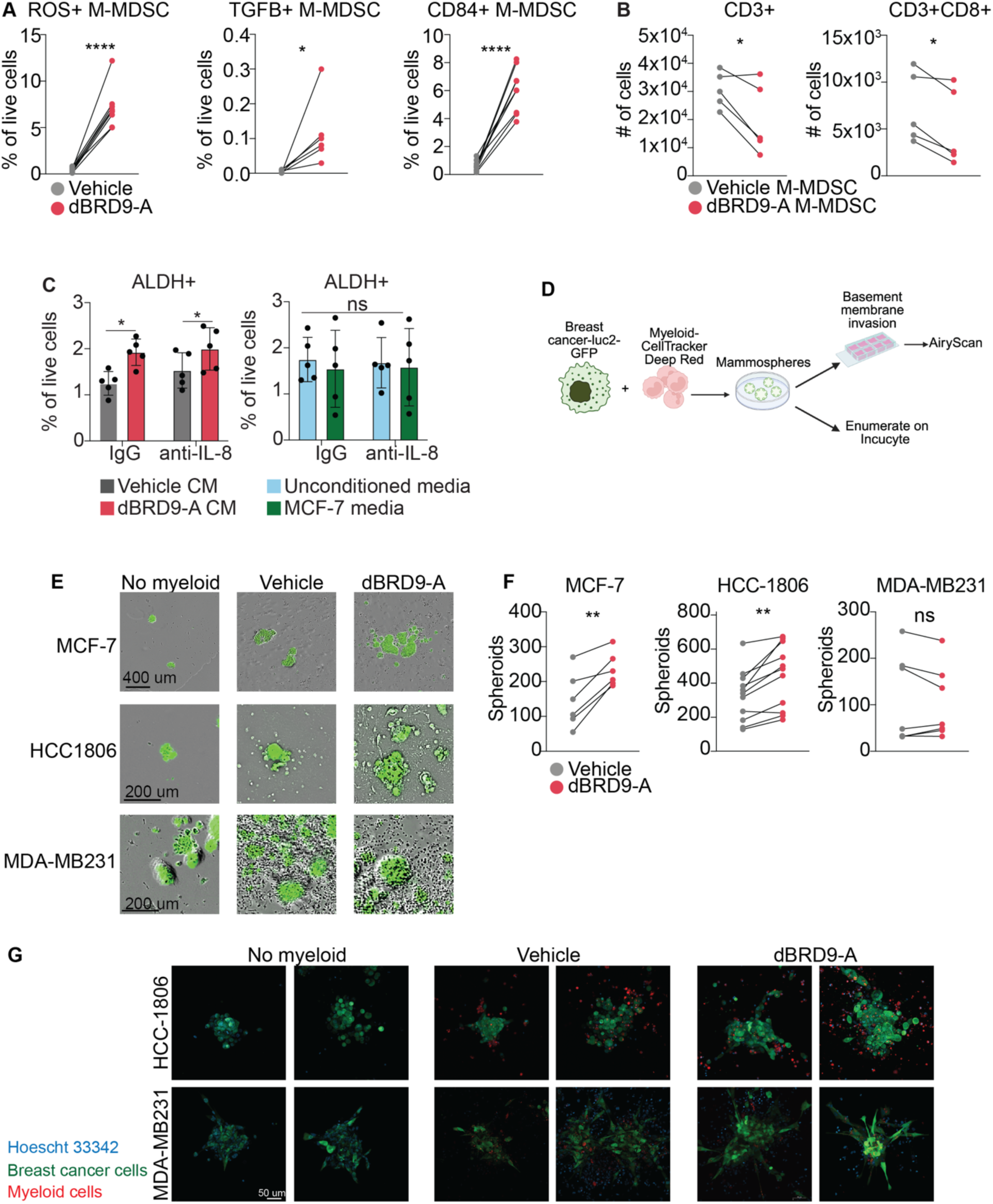
Loss of BRD9 enhances myeloid suppressor cell function. **(A)** Plots of the frequency of MDSC as percentage of all live cells that are ROS+, TGFb+, or CD84+ **(B)** Plots of the number of CD3+ T-cells (*left*) or CD3+CD8+ T-cells (*right*) after co-culture with M-MDSC that had been generated in the presence of dBRD9-A or vehicle (n=5). **(C)** Bar graphs of the percentage of ALDH+ MCF-7 cells after 72 hours of culture with dBRD9-A or vehicle conditioned myeloid media and IL-8 neutralizing antibody or IgG control (*left*) and with the MCF-7 or unconditioned myeloid media controls (*right*). **(D)** Schematic depicting the generation of mammospheres in the presence or absence of vehicle or dBRD9-A treated myeloid cells, and subsequent quantification with the Incucyte or Airy Scan microscope. **(E)** Representative Incucyte images of GFP+ mamamospheres that were generated in the absence of myeloid cells (left) or with vehicle or dBRD9-A treated myeloid cells and **(F)** summary quantification of all replicates. Each dot pair is from one biological donor. **(G)** Representative laser scan microscope images of HCC1806 (top) and MDA-MB231 mammospheres that were generated as in (F) and plated on a basement membrane overnight. Breast cancer cells stably express GFP and myeloid cells were labeled with CellTrace Deep Red. Significance calculated using paired-t-test with GraphPad Prism, p>0.05=ns, p ≤ 0.05=*, p≤0.01=**, p≤0.001=***, p≤0.0001=****.

Given these quantitative and qualitative changes in dBRD9-A MDSCs, we hypothesized that loss of BRD9 would lead to MDSCs having a stronger immunosuppressive phenotype, resulting in greater cancer aggressiveness. To test this, we first performed T cell inhibition assays. FACS-isolated MDSC were co-cultured with allogeneic T-cells, showing that despite donor-to-donor variability, M-MDSC, but not PMN-MDSC, from dBRD9-A-treated samples significantly inhibited T-cell proliferation compared to vehicle controls, confirming that these cells have stronger suppressive abilities (**Figure 7B and Supplementary Figure 8C**).

In solid tumors, MDSC contribute to immune evasion, metastasis, and aggressiveness^36^. To test whether our *in vitro*-generated MDSC may contribute to solid tumor pathogenesis, we cultured the breast cancer MCF-7 cell line in the presence of myeloid conditioned media from vehicle or dBRD9-A myeloid cells. We then assessed stemness by measuring ALDH, as MDSC have been shown to increase the stem cell properties of breast cancer cells^37^ (**Figure 7C**). Exposure of MCF-7 to dBRD9-A myeloid conditioned media significantly increased the expression of ALDH in these cells. Critically, treatment with an IL-8 neutralizing antibody showed these effects were not due to the enhanced IL-8 secretion from the BRD9 knockout cells (**Figure 7D**). In addition, co-culture of dBRD9-A myeloid cells with MCF-7 or the epithelial triple-negative breast cancer (TNBC) cell line HCC1806, led to increased formation of mammospheres *in vitro* (**Figure 7D-F**). While this increase was not observed with the more mesenchymal and claudin-low TNBC line MDA-MB231, co-culture with dBRD9-A myeloid cells did increase spheroid invasion into a basement membrane matrix in this cell line, as well as HCC1806 (**Figure 7G**). Together, these results suggest that BRD9 within the hematopoietic compartment may increase the generation of breast cancer initiating cells, although the magnitude of this effect is likely subtype specific (**Supplementary Figure 9**).

## DISCUSSION

In the hematopoietic system, prior studies have demonstrated the importance of BAF complexes in maintaining normal HSPC function^38, 39^. However, despite these reports, little was known until now about the overall binding patterns of the different complexes in human HSPC and how loss of one BAF member may affect the activity of the other complexes and therefore alter HSC function. Using a multi-omics approach, we identified chromatin binding patterns of the three main SWI/SNF complexes in adult human HSPC. Degradation of one unit, BRD9, against which both PROTAC degraders and bromodomain inhibitors are being developed, resulted in increased binding of cBAF to myeloid-associated genes, with no significant alterations in PBAF binding. This was accompanied by increased targeting of BRD4 to the CEBPA loci, a global increase in CEBPA and increased recruitment of this master regulator of myeloid cell fate to regions co-bound by cBAF. Notably, these changes were not observed with chemical inhibition of the BRD9 bromodomain.

Given the critical role that myeloid cells play within the immune system and its anti-tumoral function, we further analyzed how the myeloid compartment is impacted by its loss. Our findings show that degradation of BRD9 –but not bromodomain inhibition–contributes to enhanced differentiation of myeloid cells that secrete elevated levels of the chemoattractant chemokine IL-8. Significantly, IL-8 is elevated in many neoplasms and increased IL-8 levels are associated with poor response to immunotherapies^40, 41^, making IL-8 neutralizing antibodies a promising therapy for hematopoietic and solid tumors^42, 43^. However, our studies on these immunosuppressive cells are limited by the difficulty of characterizing MDSC by cell surface markers and by the fact that cell sorting can activate myeloid cells, making it difficult to assess their function^44^. Future studies would benefit from examining additional contexts of MDSC function, using more donor samples.

Moreover, we observed that myeloid loss of BRD9 enhanced breast cancer stemness *in vitro*. In the luminal ER+ cell we used as a model, we did not observe that this was IL-8 dependent, which is in line with MCF-7 not being responsive to IL-8^45^. Our RNA-seq data suggested that degradation of BRD9 increases levels of *CXCL1*, *CXCL6,* and *PF4* as well as the alarmin S100A8, which was confirmed by qRT-PCR. S100A8 is known to contribute to a pro-metastatic niche in breast cancer^46, 47^, however, further work will be needed to determine whether the effects of BRD9 on breast cancer are due to S100A8. Intriguingly, BRD9 is downregulated with human aging and gives rise to an MDS-like phenotype in mice^8^. However, whether the observed age-associated loss of BRD9 and increased differentiation of pro-inflammatory suppressive cells contributes to the increased incidence of solid tumors and hematological malignancies with aging remains to be determined. Together, these results suggest that BRD9 acts as a barrier to myeloid differentiation and that upon loss of BRD9, there is enhanced CEBPA activity and targeting of cBAF to myeloid genes, leading to the generation of highly suppressive, IL-8-secreting myeloid cells. Importantly, these effects are not observed with inhibition of the BRD9 bromodomain. Thus, these results suggest that complete hematopoietic loss of BRD9 may create an immunosuppressive environment permissive to the development of neoplasms.

## ACKNOWLEDGEMENTS

We thank all members of the Figueroa Lab for thoughtful discussions and suggestions. Research reported in this publication was performed in part at the Onco-Genomics Shared Resource (OGSR) and the Flow Cytometry Shared Resource (FCSR) of the Sylvester Comprehensive Cancer Center at the University of Miami, RRID: SCR_022502 and SCR_022502, which are supported by the National Cancer Institute (NCI) of the National Institutes of Health (NIH) under award number P30CA240139. The content is solely the responsibility of the authors and does not necessarily represent the official views of the NIH. E.R.A. was partially supported by the T32-CA217835 Cancer Epigenetics Training grant to the University of Miami. This work was supported by an award from the Samuel Waxman Cancer Research Foundation (SWCRF).

## AUTHOR CONTRIBUTIONS

E.R.A, and M.E.F. conceived and designed the study. E.R.A., C.R.G.S., and A.A. performed the functional experiments with myeloid and CD34+ cells and generated the ChIP-seq and ATAC-seq libraries. L.G.M. and E.R.A. performed and analyzed experiments for the in vitro co-culture with breast cancer cell lines. F.B., P.C. and R.S. assisted with the design and generation of the PRO-seq data. E.R.A. analyzed the ChIP-seq, RNA-seq, sc-RNAseq, and ATAC-seq data. O.T.P. and D.B. assisted with the MDSC T cell suppression studies. R.C assisted with the design of the mammosphere assays. L.M., D.B., R.C, and R.S. provided critical feedback. E.R.A. and M.E.F. wrote the manuscript. All authors reviewed and approved the manuscript.

## CONTACT FOR REAGENT AND RESOURCE SHARING

Further information and requests for resources and reagents should be directed to and will be fulfilled by the Lead Contact, Maria E. Figueroa.

## Data sharing statement

For all data requests, please contact.

## Statement of significance

In this study, we show that BRD9 regulates the differentiation of immunosuppressive myeloid cells in a bromodomain-independent manner and demonstrate that complete degradation of BRD9 results in increased T cell suppression and aggressiveness of breast cancer cells.

