## Supplementary material for "A ncBAF to cBAF Switch Regulates Human Hematopoietic Progenitor Myeloid Differentiation": Supplmentary figures and methods

### SUPPLEMENTARY METHODS:

#### Cell culture:

Human CD34+ hematopoietic stem and progenitor cells from mobilized PB (PB) were thawed via dropwise addition of media Iscove's Modified Dulbecco's Medium (IMDM) (Gibco, #12440-053), 20% FBS (R&D Systems, #S11150). Cells were then centrifuged for 5 minutes at 1500 RPM. The supernatant was aspirated, and cells were resuspended at a density of 1e6/mL in a stemness enhancing media consisting of IMDM with 20% human serum AB (Gemini Bio-Products, cat. 100-512), 100 ng/mL human stem cell factor (SCF) (PeproTech, #300-07), 100 ng/mL human thrombopoietin (TPO) (PeproTech, #300-018), 10 ng/mL, recombinant human Flt3-ligand (Flt3-L) (PeproTech, #300-019), 20 ng/mL human interleukin 6 (hIL-6) (PeproTech, #200-06), and 1% penicillin/streptomycin (Gibco, #15140122). Cells were plated in untreated plates and recovered at least overnight in hypoxic (4% oxygen, 5% $CO_2$ and 37 °C) conditions. Miltenyi MACS magnetic bead purification (Miltenyi, #130-100-453) was used to enrich BM MNC for CD34+ cells, which were then cultured as above. Human umbilical cord blood-derived erythroid progenitor (HUDEP) clone 2 (HUDEP-2) cell line was used as previously defined (*1*). Briefly, HUDEP-2 cells were expanded in StemSpan SFEM II (Stem Cell Technologies, #0965) supplemented with 1 uM dexamethasone (Sigma, #D4902-25MG), 50 ng/mL human stem cell factor (SCF) (PeproTech, #300-07), 3 international units (IU) IU/ml erythropoietin (EPO) (Peprotech, #100-64), 1 µg/mL doxycycline (Sigma Aldrich, #D5207-1G), and 1% penicillin/streptomycin (Gibco, #15140122) and maintained at a density of 1.5-2.5e5 cell/mL. Breast cancer cell line MCF-7 was cultured in EMEM (Quality Biologicals, #112-016-131) with 10% FBS (R&D Systems, #S11150), 1% GlutaMax (ThermoScientific, #3505006), and 0.01 mg/mL Insulin (ThermoScientific, #12585014), and 1% penicillin/streptomycin (Gibco, #15140122) in tissue culture treated plates.

#### Treatment with BRD9 chemical inhibitors:

Immediately after thawing, HSPC were expanded for 48 hours in a stemness enhancing media consisting of IMDM with 20% human serum AB (Gemini Bio-Products, cat. 100-512), 100 ng/mL human stem cell factor (SCF) (PeproTech, #300-07), 100 ng/mL human thrombopoietin (TPO) (PeproTech, #300-018), 10 ng/mL, recombinant human Flt3-ligand (Flt3-L) (PeproTech, #300-019), 20 ng/mL human interleukin 6 (hIL-6) (PeproTech, #200-06), and 1% penicillin/streptomycin (Gibco, #15140122) in hypoxia and then treated with 100 nM CFT8634 (MedChem Express, # HY-145925B), 100 nM dBRD9-A (Tocris, #6943), 1 uM I-BRD9 (Sigma-Aldrich, # SML1534), 1 uM LP99 (Tocris, #5496) or vehicle (0.01% DMSO). After treatment, HSPC were kept in culture for the given times, or used for myeloid liquid culture, with dBRD9-A and CFT8634 dosed every 48 hours and I-BRD9 and LP99 given every 24 hours.

#### Myeloid differentiation in liquid culture:

After being expanded for 48 hours and then treated with chemical inhibitors of BRD9 for 24 hours, HSPC were cultured in media promoting myeloid differentiation, containing

Iscove's Modified Dulbecco's Medium (IMDM) (Gibco, #12440-053) supplemented with 20% FBS (R&D Systems, #S11150), 100 ng/mL SCF (PeproTech, #300-07), 20 ng/mL IL-3 (PeproTech, #200-03), 10 ng/mL FLT3L (PeproTech, #300-19), 20 ng/mL GM-CSF (PeproTech, #300-23), 20 ng/mL G-CSF (PeproTech, #300-23, and 1% penicillin/streptomycin (Gibco, #15140122). Cells were cultured in normoxia at a density of 1e6/mL in untreated plates, for seven days.

**CRISPR-Cas9 knockout of BRD9:**
Immediately after thawing, HSPC were expanded for 24 hours in a stemness enhancing media consisting of IMDM with 20% human serum AB (Gemini Bio-Products, cat. 100-512), 100 ng/mL human stem cell factor (SCF) (PeproTech, #300-07), 100 ng/mL human thrombopoietin (TPO) (PeproTech, #300-018), 10 ng/mL, recombinant human Flt3-ligand (Flt3-L) (PeproTech, #300-019), 20 ng/mL human interleukin 6 (hIL-6) (PeproTech, #200-06), and 1% penicillin/streptomycin (Gibco, #15140122) in hypoxia. Prior to electroporating cells, the RNP complex was made by incubating a modified non-targeting guide (Synthego) or equimolar pool of two guides targeting BRD9 (Synthego), with recombinant Cas9 (PNA Bio, #CP02), with a ratio of 1 Cas9: 5 Guides, and 3 ug of Cas9 per reaction, for 20 minutes. Cells were then electroporated using the Neon NxT Transfection System (Thermo Scientific, #NEON1S) with 10 uL tips (Thermo Scientific, #N1096), with the conditions of 1600 V, 10 ms, and 3 pulses**.** Per reaction**,** 2e5 cells were used in a 10 uL volume. Cells were recovered in the above media for 48 hours before downstream analysis.

**Colony-forming unit assay:**
HSPC were seeded in methylcellulose, MethoCult H4435 (StemCell Technologies, #04435), in duplicate onto a 6-well SmartDish (Stemcell Technologies, #27302) at 250 cells per well per condition. For samples treated with BRD9 chemical inhibitors, CD34+ were pre-treated for 48-hours in the above stem-promoting media and then plated in methylcellulose. Chemical inhibitors were also added to the methylcellulose at the same concentration that was used in liquid culture. Colonies were scored on a STEMvision (Stemcell Technologies) after 14 days of incubation.

**Incucyte NETosis assay:**
Myeloid cells that were cultured for seven days with dBRD9-A or vehicle were resuspend in cytokine free IMDM 20% FBS media for 24 hours. Next, cells were centrifuged and resuspended in F12K media (Gibco, #21127-022) supplemented with 2% FBS and then 2e4 cells were plated per well in triplicate in a Incucyte Imagelock 96 well plate (Sartorius, #BA-04856) that was pre-coated with fibronectin (Sigma-Aldrich, #F1141-1mg). The plate was centrifuged at 50g for 1 minute to adhere the cells firmly. The plate was then placed in the Incuctye Live-Cell Analysis (Sartorius, #70667) and allowed to equilibrate at 37 °C for 30 minutes before performing the "Scan Once Now" to assess baseline viability. Then Incucyte Cytotox Green Dye (Sartorius, #4633) and Phorbol 12-myristate 13-acetate (PMA, Sigma-Aldrich, #79346) were added to final concentrations of 250 nM and 100 ng/mL, respectively. Vehicle control (PBS) was also used. The plate was then immediately placed into the Incuctye Live-Cell Analysis System and scanned with the 20X objective, with the Phase Contrast and Green channels, Standard (with lock) scan type, every 10

minutes for a total of six hours. The data was then analyzed using the Incucyte software (v2023 A Rev2, Sartorius), with Tophat and removal of dead cells based on size.

**Suppression of T-cell proliferation assay:**
Myeloid cell suppression of T-cell proliferation was performed following a modified protocol from Bruger et al (2). Lymphocytes from two different donors were isolated from healthy peripheral blood using density-based centrifugation with Lymphoprep (Stemcell Technologies, #7851), and then CD2+ cells were isolated by rosetting with sheep red blood cells (Cocalico Biologicals, # 20-1334A). The CD56+ NK cells were then removed using magnetic depletion (Miltenyi, # 130-050-401) and the T-cells frozen until future use. T-cell purity was verified by flow-cytometry, cell were stained with CD2-APC (BD, #560642), CD3-BV510 (BD, #564713), CD56-PE-Cy5 (Invitrogen, #MHCD5606), CD8-PE (BD, #555367), CD14-FITC (BioLegend, # 301804), and Sytox Blue (Thermo Fisher, #S34857). The day before the suppression assay, T-cells were thawed and cultured in complete T-cell media, consisting of RPMI supplemented with 5% Human Serum (GeminiBio, # 100-512), penicillin/streptomycin (Gibco, #15140122) and 5 U/uL IL-2 (Peprotech, # 200-02). The day of the co-culture a 96-well U-bottom tissue-culture treated plate (VWR, #10861-668) was coated with 1 ug/mL of anti-CD3 (Biolegend, # 317302) for four hours at 37 °C, and T-cells were labeled with 1 uM of CellTracker Green CMFDA Dye (Thermo Fisher, #C2925). Meanwhile, M-MDSC (CD45+, CD33+, CD14+ CD15-, HLADR-) and PMN-MDSC (CD45+, CD14-, CD15+, CD11b+, CD66b+) were FACS isolated using an BD FACS SORP Aria Fusion, from myeloid cells that had been cultured for seven days with vehicle or dBRD9-A and then stained with HLADR-PE-Cy7 (BD, #563041), CD11b-PE-Cy5 (Biolegend, #301308), CD14-FITC (Biolegend, #301804), CD15-BV605 (Biolegend, #323032), CD16-BV786 (BD, 551376), CD33-PE (BD, #555450), CD66b-biotein (Biolegend, #305120), Streptavidin-BV421 (Biolegened, #405226), CD45-AF700 (BD, #560566), and Live cell stain eFluor 780 (ThermoFisher 65-0865-14), in Brilliant Staining Buffer (BD, #566349). To assemble the co-culture, 4e4 T-cells were combined with 5.5e3 M-MDSC or PMN-MDSC in complete T-cell media with 2 ug/mL anti-CD28 antibody (Biolegend, # 302937). Cells were incubated in normoxic conditions for 96 hours. On the fourth day after co-culture, cells were incubated with FcR blocking reagent (Miltenyi, #130-050-901), followed by staining with Live/Dead eFulor 780 (eBioscience, #65-0865-14), CD3-BV510 (clone HIT3a, BD, #564713), CD8-PE (BD, #555367), CD11b-PE-Cy5 (eBioscience, #15-012-82), and CD4-BV786 (BD, #566806) in Brilliant Staining Buffer (BD, #563794). Counting beads (BD, 556288) were also including and used for enumeration of T-cells. Samples were run on a CytoFLEX Flow Cytometer (Beckman-Coulter) or FACS Canto II (BD).

**In vitro co-culture of breast cancer cell lines with conditioned media:**
Conditioned myeloid media was generated from peripheral blood CD34+ treated with dBRD9-A or vehicle, as described above. On day six of myeloid liquid culture, cells were centrifuged and resuspended in myeloid-promoting media at a density of 1e6 cells/mL, and no drug was added to the culture. On day seven, cells were centrifuged and the supernatant was collected and snap frozen. One day before co-culture, ATCC validated MCF-7 cells were seeded in 12 well plates (Costar, #3460) at 100 000 cells per well in EMEM (112-016-131, Quality Biological) supplemented with 10% FBS, 1& Pen/Strep, 1%

Glutamine and 0.01 mg/ml human recombinant insulin (Thermo Fisher Scientific, 12585014). The next morning media was replaced with myeloid media, un-conditioned media (never in contact with the myeloid cells) or MCF-7 media containing 250ng/mL of HuMax IL-8 neutralizing antibody (Invitrogen #MA5-42303), or IgG antibody (Sigma, #12511). After 72 hours, cells were collected for the identification of stem cells expressing high levels of ALDH (ALDEFLUOR™ Kit, Catalog # 01700, STEMCELL technologies). SKBR3 cells were used as positive control of the ALDH experiment. 500,000 cells per condition were incubated at 37 °C for 45 minutes. Samples were analyzed on a CytoFLEX Flow Cytometer (Beckman-Coulter).

**Mammosphere assay:**
Breast cancer cell lines were transduced with a pCDH-MSCV-HA-luc2-EF1-GFP lentivirus and GFP+ cells were FACS isolated using the Beckman Coulter CytoFLEX SRT. All myeloid cells that were used were stained with 1 uM of CellTrace Far Red (Thermo Scientific, #C34564). For mammosphere culture, 24-well plates that had been pre-coated with 120 uL of 30 mg/mL poly-HEMA (Millipore Sigma, #192066) and then dried were used. Cells were cultured at a ratio of 10 myleloid cells per one breast cancer cells, with 3,800 breast cancer cells used for HCC1806 and MCF-7, and 7,900 used for MDA-MB231. Cells were plated together in media consisting of Advanced DMEM/F12, without phenol red (Thermo Scientific, #241041025) with 20 ng/uL Animal-Free Human EGF (Peprotech, #AF-100-15), 20 ng/uL bFGF (Thermo Scientific, #100-18B-50ug), 2% GlutaMax (Gibco, #35050-061), 1% B27 supplement (Life technologies, #17504044), and 1% Penicillin-Streptomycin (Thermo Scientific, #15140122). After seven days of culture, mammospheres were enumerated using the Incucyte Sx5. For imaging, mammospheres were transferred 24-hours ahead of time, using BSA coated tips to a μ-Slide 8 well high (Ibidi, #80827) slide that was coated in 50 uL of Geltrex (Gibco, #A40000467-02). One hour prior to imaging, Hoescht 33442 (Thermo Scientific, #62249) dye was added to a final concentration of 1.5 uM. Live imaging of the spheres was performed using a Zeiss LSM 980 with AiryScan2 microscope, using the 20X objective, with 1X zoom, with MPLX SR-8Y mode, averaging, and bidirectional scanning. The Airyscan Processing function was then used on the z-stack.

**Isoplexis Single-cell Secretome:**
Myeloid cells that were cultured for seven days with dBRD9-A or vehicle were resuspend in cytokine free IMDM media supplemented with 10% FBS, 1% GlutaMax (ThermoScientific, #3505006), and 1% penicillin/streptomycin (Gibco, #15140122) for 24 hours. One hour before running the Isoplexis, the Single Cell Inflammation Panel, (Isoplexis, #PANEL-1003-4) chip was thawed at room temperature. Cells were then washed in PBS and resuspended in 100 uL of Isoplexis Stain Master Mix, which was previously resuspended in DMSO and diluted in PBS. After a five-minute incubation in the dark, 500 uL of IMDM 10% FBS, 1% Glutamax and 1% penicillin/streptomycin was added and the cells were incubated for 10 minutes at 37 °C. Next the cells were counted and then centrifuged and resuspended at a density of 7.5e5 cells/mL in IMDM 10% FBS, 1% Glutamax and 1% penicillin/streptomycin. In the tissue culture hood, 40 uL of the sample was loaded into the chip, which was immediately run on the Isolight (Isoplexis,s #1000-

1) with the single-cell program. Data was analyzed using the IsoSpeak v2.9.0.33 software (Isoplexis).

**Western blot of nuclear fractions:**
To extract nuclear protein, cells were resuspended in buffer containing 25 mM HEPES pH 7.6, 5 mM MgCl2, 25 mM KCl, 0.05 mM EDTA, 10% glycerol and 0.1% NP40, 1X protease inhibitor (Sigma-Aldrich, #11836170001), 1 mM DTT and 1 mM phenylmethylsulfonyl fluoride (PMSF) at a density of 10 million cells/mL and incubated on ice for 10 minutes. Sample was then centrifuged for 10 minutes at 3000 RPM at 4 °C. Cytoplasmic supernatant was discarded and nuclear pellet was resuspended in 150 mM NaCl, 20mM Hepes pH 7.9, 1% NP40, 1X protease inhibitor, 10% glycerol, 1 mM DTT and 1 mM PMSF at a density of 0.1 mL per 10 million starting cells and incubated on ice for 10 minutes. Samples were then sonicated in Biorupter (Diagenode, Bioruptor Plus) for 5 cycles of 30 seconds On/30 seconds Off. Samples were then centrifuged at 10,000 RPM for 10 minutes at 4 °C. Protein was quantified using the Bradford assay (Bio-Rad, #5000006) according to the manufacturer's instructions. Samples were diluted to equivalent concentrations using LDS loading buffer (Thermo-Scientific, #NP0007) with 5% β-mercaptoethanol (MP Biomedicals, #190242) and denatured for 5 minutes at 95°C for 5 minutes. Samples were loaded into a 4-12% Bis-Tris gel (Thermo Scientific, #NP0322BOX or # NP0315BOX) and ran with 1X MES-SDS running buffer (Thermo Scientific, #NP0002) and the Chameleon Ladder (LICor-Biosciences, #928-60000) was also used. Protein was transferred to a nitrocellulose membrane (Bio-Rad, # 1704158) using the Bio-Rad Turbo transfer and turbo transfer buffer (Bio-Rad, #10026938) with the Mixed MW 1.3A-25W program. Membranes were blocked for one hour at room temperature in Intercept PBS blocking buffer (LI-COR, # 927-70001. Then membranes were incubated with primary antibody diluted in TBS-Tween 5% milk at 4°C overnight, followed by secondary antibody in TBS-Tween 5% milk at room temperature for one hour. Antibodies used were as follows: rabbit anti-BRD9 (CST, #48306, 1:5000 dilution), mouse anti-HDAC1 (CST, #5356, 1:2000 dilution), rabbit anti-CEBPA (Diagenode, #C15410225-100, 1:1000 dilution), rabbit anti-CEBPB (CST, #43095S, 1:1000), rabbit anti-CEBPE (Thermo-Scientific, #PA5-99151, 1:1000), rabbit anti-H3 (Abcam, #ab1791, 1:3000 dilution), and mouse anti-H3 (Abcam, # ab10799, 1:3000 dilution). Fluorescence detection was performed using the secondary antibodies anti-Rabbit IR-Dye 800 CW (LI-COR, #92632213) and anti-Mouse IR-Dye 680 RD (LI-COR, # 925-68070). Densitometry was performed using the LI-COR Image Studio software. For all samples, intensity of protein of interest was normalized to HDAC1 loading control.

**Western blot of chromatin fractions:**
For analysis BRD4 in the chromatin fraction, protein was extracted using the Pierce Subcellular Protein Fractionation Kit (Thermo Scientific, #378840), using half of the recommended volume of buffers, and quantified using the Pierce BCA Protein Assay (Thermo Scientific, #23227). Electrophoresis was performed as above, using NuPAGE 7% Tris-Acetate Protein Gel (Thermo Scientific, #LA0050), Tris Acetate SDS Buffer (Thermo Scientific, EA0358BOX), and HiMark high MW protein standard (Thermo Scientific, #LC5699). For the transfer and blotting were performed as above, with the exception of using the Standard "SD" program, 25V-1.0A. Antibodies were as follow,

BRD9 (CST, #48306S, 1:5000), (BRD4 (Bethyl, #A700-004, 1:1000), and HDAC1 (CST, #53565, 1:1000 dilution). Fluorescence detection and quantification was performed as described above.

**Reverse-transcription PCR and quantitative PCR:**
RNA was extracted using the RNAeasy Plus Micro Kit (Qiagen, #74034) according to the manufacturer's instructions and reverse transcribed using the Verso cDNA Synthesis Kit (Thermo Fisher Scientific, #AB1453) according to the manufacturer's recommendations. Quantitative reverse-transcription PCR was performed with Fast SYBR Green Master Mix (Thermo Fisher Scientific, # 4385617) and the QuantStuido 5 (Thermo Fisher Scientific). The reactions were performed in triplicate for each primer. Relative mRNA expression was normalized to GAPDH as an endogenous control using the 2-DDCT method. For nascent qRT-PCR of MPO, cDNA was RNAse (Millipore-Sigma, # 3335399001) treated prior to qPCR and the primer was designed to span an intron-exon boundary. A reverse transcriptase free reaction was also performed as a control for the nascent qRT-PCR samples.

**IL-8 Immunoassay of conditioned media:**
Immunoassay to quantify secreted IL-8 in conditioned media from vehicle or dBRD9-A-treated myeloid cells was performed using the ProQuantum™ Human IL-8 Immunoassay Kit (Invitrogen, #A35575) following the manufacturer's instructions. To generate the myeloid media, myeloid cells were cultured for seven days in myeloid-promoted media and then centrifuged and resuspended in cytokine-free Iscove's Modified Dulbecco's Medium (IMDM) (Gibco, #12440-053), 20% FBS (R&D Systems, #S11150), penicillin/streptomycin (Gibco, #15140122). Cells were cultured for 24 hours, and then the media was collected and snap frozen. The assay was run using a QuantStudio 5 (Thermo Fisher Scientific).

**Flow-cytometry of myeloid and MDSC associated marks:**
Analysis of human myeloid cell surface markers was performed on the seventh day of culture in myeloid promoting media. Samples were stained with CD34-APC (BD, #340441), CD11b-PE-Cy5 (Biolegend, #301308), CD15-BV605 (Biolegened, #323032), CD33-PE (BD, #555450), CD14-FITC (Biolegened, #301804), HLA-DR-PE-Cy7 (BD, #563041), CD16-BV786 (BD, #551376), and CD66b-biotin (BD, #567941), Viability-eFluor-450 (eBioscience, #65-0863-14), and Streptavidin-APC-Cy7 (BD, #554063) in Brilliant Staining Buffer (BD, #566349). Samples were fixed in 4% paraformaldehyde (Electron Microscopy Sciences, #15710) and analyzed on an LSR-Fortessa-HTS (BD).

**Flow-cytometry of NCP and GMP associated markers:**
After being treated for 48 hours with chemical inhibitors of BRD9 or vehicle control, HSPC were stained with CD1c-BV510 (Biolegend, #312220), CD14-BV510 (Biolegend, #367124), CD16-BV510 (Biolegend #302048), CD19-BV510 (BD, #562953), CD56-BV510 (BD, #562953), CD141-BV510 (BD, #563298), CD11b-PerCP-Cy5.5 (Biolegend, #393106), CD15 -BV605 (BD, #323032), CD34-APC (BD, #340441), CD45RA-FITC, (Invitrogen, #MHCD45RA01), CD64-PE (Biolegend, #305008), CD66b-BV421 (BD, #562940), CD45-PE-CF594 (BD, #562279), CD115-PE-Cy7 (Biolegend, #347308),

CD123-BV711 (BD, #568228), and Viability-LIVE/DEAD Fixable Blue Dead Cell (ThermoFisher, #L34962) in Brilliant Staining Buffer (BD, #566349). Samples were fixed in 4% paraformaldehyde (Electron Microscopy Sciences, #15710) and analyzed on an LSR-Fortessa-HTS (BD).

**Intracellular staining of IL-8 for flow-cytometry analysis:**
Intracellular staining was performed on myeloid cells that were cultured for seven days with inhibitors of BRD9 or vehicle controls. To prevent secretion of IL-8, cells were treated with 2 uM GolgiPlug (Brefeldin A, BD, #555028) for four hours in a 5% CO2 and 37 °C tissue-culture incubator. During this period, half of the cells were also stimulated with a combination of 100 ng/mL LPS (Sigma-Aldrich L2880-100MG), 100 ng/mL PMA (Sigma-Aldrich, # 79346-1MG), and 1 ug/mL Ionomycin (Sigma-Aldrich #I0634-1MG) as a positive control. Cells were then centrifuged and resuspended in diluted FcR block (Miltenyi, #130-050-901) and incubated for five minutes at room temperature, followed by staining for the cell surface markers CD11b-PE-Cy5 (Biolegend, #301308), CD15-BV605 (BD, #323032), CD33-PE (BD, #555450), CD14-FITC (Biolegend, #301804), HLA-DR-PE-Cy7 (BD, #563041), CD16-BV786 (BD, #551376), CD66b-biotin (BD, #567941), Streptavidin-APC-Cy7 (BD, #554063) and then viability staining with Viability-eFluor-450 (eBioscience, #65-0863-14). Cells were then fixed in 4% paraformaldehyde (Electron Microscopy Sciences, #15710) for 15 minutes at room temperature and then stored overnight at 4 °C. The following day, intracellular staining was performed using IL-8-APC (Thermo Scientific, #17-8088-42. 5:100 dilution) diluted in 1X BD Perm/Wash buffer (BD, #554715). After staining cells were washed twice using in 1X BD Perm/Wash buffer and then analyzed on an LSR-Fortessa-HTS (BD).

**Intracellular staining of TGFb:**
Intracellular staining of TGFb was performed on myeloid cells that were cultured for seven days with inhibitors of BRD9 or vehicle controls. To prevent secretion of TGFb, cells were treated with 2 uM GolgiStop (Monensin, BD, #554724) for four hours in a 5% CO2 and 37 °C tissue-culture incubator. Cells were then centrifuged and resuspended in diluted FcR block (Miltenyi, #130-050-901) and incubated for five minutes at room temperature, followed by staining for the cell surface markers CD14-BB70 (BDBiosciences, #568525), CD49d-RB780 (BDBiosciences, #755833), CD66b-AF700 (BioLegend, #305114), CD15-BUV805 (BDBiosciences, #568935), CD11b-BV605 (BioLegend, #101257), CD16-BV421 (BDBiosciences, #562874), CD33-BV711 (BioLegend, #303424), LOX1-BV786 (BDBiosciences, #772187), CD10-PE-Cy5 (eBiosciences, #772187), CD84-PE (BioLegend, #326008), HLA-DR-PE-Cy7 (BioLegend, #327018) diluted in Brilliant Staining Buffer (BD, #566349), for 25 minutes on ice. Then viability staining with eFluor780 Fixable Viability Dye (ThermoScientific, # 65-0865-14) was performed. Cells were then fixed in 4% paraformaldehyde (Electron Microscopy Sciences, #15710) for 15 minutes at room temperature and then stored overnight at 4 °C. The following day, intracellular staining was performed using TGFB1-PE-CD594 (Thermo Scientific, #17-8088-42, 5:100 dilution) diluted in 1X BD Perm/Wash buffer (BD, # 562422). After staining cells were washed twice using in 1X BD Perm/Wash buffer and then analyzed on an Cytek Aurora analyzer.

**CellROX Staining:**
Staining for ROS and MDSC markes was performed on myeloid cells that were cultured for seven days with inhibitors of BRD9 or vehicle controls. Cells were stained for surface markers CD14-BB70 (BDBiosciences, #568525), CD49d-RB780 (BDBiosciences, #755833), CD66b-AF700 (BioLegend, #305114), CD15-BUV805 (BDBiosciences, #568935), CD11b-BV605 (BioLegend, #101257), CD16-BV421 (BDBiosciences, #562874), CD33-BV711 (BioLegend, #303424), LOX1-BV786 (BDBiosciences, #772187), CD10-PE-Cy5 (eBiosciences, #772187), CD84-PE (BioLegend, #326008), HLA-DR-PE-Cy7 (BioLegend, #327018), and CD34-APC (BDBiosciences, #340441) diluted in Brilliant Staining Buffer (BD, #566349), for 25 minutes on ice, followed by viability staining with eFluor780 Fixable Viability Dye (ThermoScientific, # 65-0865-14). Cells were then incubated in freshly prepared 5 uM CellROX reagent (ThermoFisher, #C10492) for 30 minutes at 37 °C. After washing, samples were immediately analyzed on the Cytek Aurora Cell Sorter.

**ChIP-seq of human HSPC:**
ChIP-seq of HSPC was performed in mobilized peripheral blood CD34+ cells that were expanded in hypoxic conditions in stemness promoting media consisting of IMDM with 20% human serum AB (Gemini Bio-Products, cat. 100-512), 100 ng/mL human stem cell factor (SCF) (PeproTech, #300-07), 100 ng/mL human thrombopoietin (TPO) (PeproTech, #300-018), 100 ng/mL, 10 ng/mL recombinant human Flt3-ligand (Flt3-L) (PeproTech, #300-019), 20 ng/mL human interleukin 6 (hIL-6) (PeproTech, #200-06), and 1% penicillin/streptomycin (Gibco, #15140122) for 48 hours. Cells from each donor (n=2 per protein examined) were then split in half and treated with 100 ng/mL dBRD9-A or vehicle control for 72 hours, while still in hypoxia. Samples treated with vehicle were used for the baseline analysis. ChIP was performed with the Ideal TF ChIP Kit (Diagenode, # C01010055) following the manufacturer's instructions with the following modifications. Briefly, 5 million cells per IP were used. Double crosslinking was performed by adding 16 uL of Cross-link Gold (Diagenode, #C01019027) per 5 mL of cells and then incubating at room temperature for 30 minutes with agitation. Samples were then washed twice using PBS and then formaldehyde (Sigma-Aldrich, #252549) crosslinking was performed by fixing the cells for 10 minutes at room temperature in a 1% formaldehyde solution. The volumes of cell lysis and shearing buffers were adjusted to use 0.5 mL of 1L1b buffer and 0.15 mL of i2L2b buffer per one million cells, along with 250 uL of shearing buffer per 10 million cells. For BRD9, BRD4, and CTCF immunoprecipitations, Drosophila spike-ins (Active Motif, # 53083) were added, following the manufacturer's instructions. Samples were sonicated using the Diagenode Bioruptor Pico in 1.5 mL Pico tubes (Diagenode, #C30010016) for eight cycles of 30 seconds ON/30 seconds OFF. Immunoprecipitation was performed for 16 hours using the following antibodies: ARID1A (CST, #12354S, 6 uL/IP), ARID2 (CST, #82342S, 7.5 uL/IP), BRD4 (Bethyl, # A700-004, 0.5 ug/IP), BRD7 (CST, #15125S, 7.5 uL/IP), BRD9 (Bethyl, #A700-153, 0.5 ug/IP), BRG1 (Epicypher, #13-2002, 1 ug/IP), CEBPA (Diagenode, #C15410225-100, 5 ug/IP), CTCF (Diagenode, #K08121004, 2 ug/IP), DPF2 (Abcam, #ab134942, 5 ug/IP), GLTSCR1 (CST, #45441S, 12.5 uL/IP), and H2Av spike-in (Active Motif, #61686). For the final DNA cleanup, the Qiagen minElute kit (Qiagen, #28004) was used. Sample concentration was measured using the Qubit DNA high sensitivity kit (Thermo Fisher Scientific, #Q32851) and

enrichment over input was performed using qPCR with SYBR green (Thermo Fisher Scientific, #4385617). Libraries were then prepared using the CUTANA Library Prep Kit (Epicypher #14-1001 and #14-1002) following the manufacturer's instructions with the following modifications. Up to 10 ng of DNA was used for library prep and all samples were amplified for 16 cycles and for the final Ampure bead cleanup, a ratio of 0.85X was used. Prior to sequencing, libraries were quantified and pooled together and another Ampure cleanup was performed to remove any remaining indices. Libraries were sequenced by the University of Miami Onco-Genomics Shared Resource using the NovaSeq and NovaSeq X plus, with 100-150 bp single- or paired-end reads.

**ChIP-seq data processing:**
FastQC was used to evaluate library quality (*3*). Illumina adapters were trimmed and fragments less than 25 bp were removed using Cutadapt (v2.6) (*4*). All samples were analyzed as single-end and only Read 1 was used. Reads were aligned to hg38 using Bowtie2 (v2.2.5) or a merged hg38 and dm6 genome if they contained Drosophila spike-ins (*5*). Duplicate reads were flagged using Picard (v2.21.2) with the picard MarkDuplicates I = aligned_sort_name O = aligned_dup_name M = marked_dup_metrics ASSUME_SORTED = true REMOVE_DUPLICATES = false VALIDATION_STRINGENCY = LENIENT (*6*). Duplicate and multimapped reads were then removed with samtools (v1.9) with the commands: samtools view -b -F 4 -q 5 and samtools view -b -F 1024 -L (*7*). For each individual sample, bigwig files for visualization were generated using deeptools (v3.3.1) bamCoverage with a binSize=5 and CPM normalization (*8*). Aligned bam files for each protein type for each condition were also pooled together and then bigwigs for the $\log_2$(pooled IP/pooled Input) were generated using deeptools bamCompare with the --operation log2 and --scaleFactorsMethod readCount. In addition, bigwig files were prepared using the same command for $\log_2$(dBRD9-A/vehicle) or $\log_2$(I-BRD9/vehicle). Peak calling for individual IPs was performed using the callpeak function from the Model Based Analysis of ChIP-seq 2 (MACS2 v.2.2.4) program, specifiying macs2 callpeak -t IP_bam -c Input_bam -f BAM -B -n name -g hs --nomodel -q 0.001 --SPMR --keep-dup all --extsize 200 (*9*).

**Identification of BAF complex peaks and differential peaks:**
Peak calling and differentially peak calling for each protein assessed was performed using csaw (v1.24.3) using the uniquely mapped reads for each sample and its corresponding input, as well as the "narrowPeak" file generated by macs2. For each protein, the correlateReads function was used to compute the average fragment length. The windowCounts function was then used to count reads into windows (width=150 for all BAF proteins and width=10 for CTCF and CEBPA), and low abundance windows were filtered out using the filterWindowsGlobal on binned data (width=15000 for ARID1A, ARID2, BRD7, BRD9, DPF2, and GLTSCR1, width=2000 for BRG1 and width=10000 for CEBPA and CTCF), with a minimum fold-change of 3 for BRD9 and 5 for all other proteins assessed. For BRD9, normalization to spike-ins was performed using normFactors and the spike-in counts. For all other proteins, normalization for composition bias was performed using normFactors and then data was converted to an edgeR object. Windows less than 100 bp apart were clustered into regions and a maximum cluster width of 5 kb was set. Differential peaks for each protein were defined as only those with FDR < 0.05.

To define baseline peaks for each protein, windows were filtered to keep those that had at least CPM >1 for each vehicle sample. Given the background for the BRD9 antibody, baseline peaks were defined as only those that had a significant decrease (FDR < 0.05) with dBRD9-A treatment. Bedtools merge (v2.29.0) was then used to define baseline cBAF as regions with ARID1A or DPF2 vehicle peaks, ncBAF as regions with GLTSCR1 or BRD9, and PBAF as regions with BRD7 or ARID2 (*10, 11*). To define differential BAF binding, we defined differential ncBAF peaks as those that had significant (csaw FDR < 0.05) changes in BRD9 or GLTSCR1 and differential cBAF peaks that had changes in ARID1A or DPF2. Differential "core" peaks were defined as those peaks with significant changes in BRG1, that had no significant change in any of the other BAF complex proteins assessed.

**Functional and genomic annotation of ChIP-seq data:**
Significant (FDR < 0.05) differential peaks were annotated to the nearest transcription start site of RefSeq hg38 using the R package ChIP-enrich (v2.14.0) (*12*). Genomic annotation of peaks was performed using the R-package Genomation (v1.22.0) (*13*), to hg38 genomic features or active and poised enhancers previously identified in young HSCe (*14*). Heatmaps of baseline and differential peaks were plotted using the $log_2$(IP/Input) or $log_2$(Treatment/Vehicle) bigwig files generated using deepTools bamCompare and plotHeatmap functions (*8*). Peaks were visualized using the UCSC genome browser. Transcription factor motif analysis was performed using the Homer findMotifsGenome function (version 4.10-0) (*15*). Heatmaps of the $log_{10}$(p-value) transcription factor enrichment were plotted using ComplexHeatmap (version 2.6.2) (*16*).

**OMNI-ATAC-seq of HSPC:**
ATAC-seq of HSPC treated for 72 hours with vehicle or dBRD9-A was performed following the OMNI-ATAC-seq protocol (*17*). Vehicle treated samples were used for baseline analysis of accessibility in HSPC. Briefly, 60,000 cells per sample were used as input and the optional DNAse (Thermo Scientific, #AM2222) treatment was performed. Cells were then lysed in ice-cold ATAC-RSB buffer, consisting of 10 mM Tris-HCl, pH 7.5, 10 mM NaCl, and 3 mM $MgCl_2$, supplemented with 0.1% NP40, 0.1% Tween-20, and 0.01% digitonin. Nuclei were then washed in ATAC-RSB with 0.1% Tween-20. Transposition was performed by incubating nuclei TDE1 TD enzyme (Illumina, #20034210) for 30 minutes at 37 °C. DNA was then purified using the DNA Clean and Concentrator kit (Zymo, #D4014). Barcoding of transposed fragments was performed by using dual indices (custom from IDT) with NEBNext Ultra II Q5 2X Master Mix (NEB, #M0544S). Libraries were sequenced by the University of Miami Onco-Genomics Shared Resource using the NovaSeq with paired-end 150 bp sequencing.

**OMNI-ATAC-seq data processing:**
FastQC was used to evaluate library quality (*3*). Using Cutadapt (v2.6), Illumina adapters were trimmed, fragments less than 25 basepairs were removed, and reads were trimmed to 75 base-pairs, to allow for better detection of nucleosome free regions, using the command cutadapt -a CTGTCTCTTATA -A CTGTCTCTTATA -m 25 --length 75 -o name_R1_trim.fastq.gz -p name_R2_trim.fastq.gz fastq_R1_fullpath fastq_R2_fullpath (*4*). Data was aligned to hg38 using Bowtie2 (v2.2.5) using the command: bowtie2 -p 20

--mm --local --very-sensitive-local --no-unal --no-mixed --no-discordant -I 10 -X 1000 -x bwt2_idx -1 trim_R1_fullpath -2 trim_R2_fullpath | samtools view -Sb - > aligned_name) 2>name.log (*5*). Aligned files were then coordinate sorted and indexed with samtools (v1.9) and used to generate a bam file with mitochondrial reads removed. Samtools was then used to remove unmapped, mate unmapped, not primary alignment, mitochondrial reads, and orphan reads. Duplicate reads were flagged with Picard (v2.21.2) (*6*) with the command: picard MarkDuplicates INPUT=FILT_BAM_FILE OUTPUT = TMP_FILT_BAM_FILE METRICS_FILE= DUP_FILE_QC VALIDATION_STRINGENCY = LENIENT ASSUME_SORTED = true REMOVE_DUPLICATES = false; mv TMP_FILT_BAM_FILE FILT_BAM_FILE. Duplicate reads were then removed with samtools: samtools view -F 1804 -f 2 -b FILT_BAM_FILE > FINAL_BAM_FILE; samtools index FINAL_BAM_FILE. Quality control of the library complexity, TSS score, fraction of mitochondrial reads, and fragment length was performed using the R (v4.0.2) package ATACseqQC (v1.14.4) (*18, 19*). The deduplicated, mitochondrial free bam files were then used as input to the Encode ATAC-seq Pipeline, which generated peak and bigwig files (*20*).

**Analysis of differentially accessible regions (DAR):**
Before performing DAR analysis, iterative peak calling was used to identify a merged peak set for the vehicle and dBRD9-A samples, with an extension of 250, a scorePerMillion cutoff of 1, and a selectionRule of (n+1)/2 (*21*). FeatureCounts (v1.6.4) was used to generate a matrix of counts for the consensus peak set, using the command: featureCounts -p -F GTF -T 15 -t "peak" -g "peak_id" -a consensus_peaks_gtf -o output_name --fracOverlap 0.2 bam_list (*22*). Counts were then normalized for GC content using EDAseq with the function: dataOffset = withinLaneNormalization( reads.peak.m, y = gcContentPeaks, num.bins = 20, which= "full", offset = FALSE) (*23, 24*). DAR analysis was then performed using DESeq2, specifying fitType="local" and using a varianceStabilizingTransformation (VST) of the counts, along with a design that took into account the donor identification (*25*). Shrinkage of counts was performed using the associated ashr method (*26*). Significant regions were defined as those with a p-adjusted < 0.05 and an absolute $\log_2$FoldChange > 1.5. Annotation to the nearest transcription start site as well as gene-ontology processes was performed with the R package chipenrich (version 2.14.0) (*27*). Motif analysis was performed using HOMER (v4.10-0).

**RNA-seq of myeloid cells:**
RNA was extracted from myeloid cells that were treated with BRD9 inhibitors or vehicle control for seven days (n=4 donors) using the RNAeasy Plus Micro Kit (Qiagen, #74034) according to manufacturer's instructions. The ThermoFisher Scientific DNA-free™ DNA Removal Kit (#AM1906) was used to remove any trace amounts of DNA prior to library preparation. First-stranded libraries with ERCC spike-in controls (ThermoFisher Scientific, #4456740) were prepared by the University of Miami Onco-genomics Sequencing Core using the Kapa HyperPrep kit (#0810595200) with Kapa RiboErase (#07962347001). Libraries were sequenced on the NovaSeq with 100 bp paired-end sequencing.

**RNA-seq alignment:**
Using Cutadapt (v2.6), adapters were removed from both reads and reads less than 25 basepairs were discarded (*4*). Reads were aligned to the hg38 gencode v43 reference genome using the STAR aligner (v2.7.3), specifying the following parameters: STAR --runThreadN 5 --genomeDir genome_path --readFilesIn trim_R1_fullpath trim_R2_fullpath --outFileNamePrefix name_ --outFilterType BySJout --outFilterMultimapNmax 20 --alignSJoverhangMin 8 --alignSJDBoverhangMin 1 --outFilterMismatchNmax 999 --alignIntronMin 20 --alignIntronMax 1000000 --alignMatesGapMax 1000000 --readFilesCommand gunzip -c --outSAMtype BAM SortedByCoordinate --outWigType bedGraph --outWigStrand Stranded --outWigNorm RPM --alignEndsType EndToEnd (*28*).

**Differential gene-expression analysis:**
Gene counts were calculated using FeatureCounts (v1.6.4) with the hg38 genecode annotation file that also contained the ERCC spike-ins, using the command: featureCounts -T 10 -a gtf_file --minOverlap 1 -s 2 -o name_counts.txt sorted_bam (*22*). Differential gene expression analysis was performed using the package DESeq2 (v1.30.0) (*29*). A multifactor design was used to control for effects of donor variation, while measuring the effect of treatment. Size factors were estimated using the ERCC spike-in controls and then dispersions were calculated using samples from all treatment groups. For differential analysis, spike-ins were removed, and contrasts were established for pair-wise comparisons of each treatment versus vehicle. The log-fold change was corrected using the ashr method (*26*). Significant genes were defined as having a fold change >1.5 and p-adjusted <0.05. Regularized log-counts (rld) and the Wald-statistic were generated with DESeq2 and genes were annotated to ensembl version 109. For GSEA (v4.3.2) the Wald statistic ranked list was used with the c2.all.v2023.2.Hs.symbols gene set using the weighted enrichment score (*30*). For the heatmap of genes that are differentially expressed with treatment of any of the BRD9 inhibitors, normalized counts were plotted with ComplexHeatmap (v2.6.2) using distance correlation and the average clustering method (*16*). To determine which cell types were associated with up- or down-regulated genes, EnrichR was run on the list of significant gene names and the PanglaoDB dataset was used for cell type enrichment (*31*).

**Single-cell RNA-seq:**
Single-cell RNA-seq (sc-RNA-seq) was performed on human myeloid cells that were treated with dBRD9-A or vehicle for seven days, as described above (n=2 biological donors). Prior to library preparation, quality control for sample viability and concentration was performed using the Nexcelom Cellometer K2 with ViaStain APOI Staining Solution. Libraries were then prepared by University of Miami Onco-Genomics Shared Resource using the 10X Genomics 3' v3.1 single cell RNA-Seq Kit and sequenced on a NovaSeq with paired-end 100 bp sequencing, targeting 10,000 cells and a depth of ~440 M reads per sample. Raw sequencing data was demultiplexed and converted to the FASTQ files by executing cellranger mkfastq (10x genomics).

**Single-cell RNA-seq processing and annotation:**

Alignment to hg38 (Gencode v43) and generation of a filtered gene count mtx matrix that contained only those barcodes associated with cells was generated using the nfcore scrnaseq pipeline (v2.3.2) with the command: nextflow run nf-core/scrnaseq --input config.csv -profile docker --outdir /my_outdir --aligner cellranger -r 2.3.2 --fasta genome.fa --gtf annotation.gtf --genome GRCh38 -w /working_dir (*32*). The matrix was then converted to h5ad format and quality control filtering was performed using the Median Absolute Deviation (MAD) method to remove cells that differed by five MADs or had greater than eight percent percent mitochondrial reads (*33*). The SoupX package (v1.6.2 ) was then used to remove ambient RNA (*34*). Doublet detection and removal was performed using the R package scDblFinder. (v.1.12.0) (*35*). Before filtering, we had a median of 9,405 cells, after all filtering steps, there was a median of 7,921 cells. Normalization of the SoupX corrected count matrix was then performed across all samples (both vehicle and dBRD9-A) using the R package Seurat (v4.3.0.1) NormalizeData function (*36*). Using the Seurat splitObject command the vehicle and dBRD9-A conditions were then split and SCTransform was run on each condition, specifying vst.flavor = "v2". Then data integration was performed using the Seurat functions SelectIntegrationFeatures with nfeatures=3000, PrepSCTIntegration, FindIntegrationAnchors with normalization.method = "SCT", and IntegrateData with normalization.method = "SCT". After integration, there were 27,215 cells with 46,448 features (genes). Principal component analysis of the integrated data was performed using the RunPCA function and uniform manifold approximation projection was run using the RunUMAP function with dims = 1:40. Clustering was then done usine the FindNeighbors function with dims = 1:40, followed by FindClusters with resolution of 0.3. UMAP plots were generated on data that was then normalized using the NormalizeData function and plotted with the FeaturePlot function. For plotting purposes, populations with less than 50 cells were omitted. Cell annotation was performed with the R packages celldex (v1.8.0) and SingleR (v2.0.0) with a combined reference of the MonacoImmuneData and HumanPrimaryCellAtlasData data sets and label.main for each (*37*). Poor quality and ambiguous assignments were removed with the pruneScores function. The gene count matrix was then annotated to MDSC as defined by the general PanglaoDB gene signature, or the primary breast cancer associated MDSC identified by Alshetaiwi et al. using the R package AUCell and the function AUCell_run (*38, 39*). Histograms for each dataset were evaluated using the AUCell_plotHist function and the thresholds for each annotation were set to 0.2. The SingleR cell annotations were then overwritten, if necessary, with the MDSC annotation. The MDSC marker genes were identified using the Seurat function FindConservedMarkers, with min.cells.group = 50 and a heatmap was then plotted with DoHeatmap.

**Single-cell RNA-seq differential gene expression and abundance analysis:**
For differential analysis, the integrated matrix was converted to a SCE object using the Seurat as.SingleCellExperiment function. Differential abundance analysis was then performed using edgeR (v3.40.0), with a blocking factor for donor ID, and running the estimateDisp function on the abundances and specifying trend = "none". To test for difference, the glmQLFTes function was used (*40*). For pseudobulk differential expression analysis between the vehicle and dBRD9-A for each cell type, DESeq2 (v.1.42.0) was used. Briefly, the counts were aggregated based on cell type and treatment groups using

the aggregate.Matrix function with fun = “sum”. A dds object was then generated by running DESeqDataSetFromMatrix(round(cluster_counts), colData = cluster_metadata, design = ~ condition) and differential expression analysis was performed using the DESeq function, followed by lfcShrink. Significant genes were defined as those with a p-adjusted <0.05. To visualize differentially expressed genes, normalized counts were extracted from the dds object using counts(dds_obj, normalized = TRUE) and then a heatmap was plotted using the R package pheatmap.

**PRO-seq library preparation:**
PRO-seq was performed on HSPC (n=3 biological replicates) that were thawed and recovered for 48 hours, as described above, and the treated for three hours with 100 nM of dBRD9-A or vehicle control. Library preparation was carried out following established protocols from previous studies (*41-43*). In brief, nuclear run-on assays were performed using two million nuclei, which were mixed with spike-in Drosophila S2 cell nuclei. The mixture was incubated for 3 minutes at 30°C in the presence of 25 µM Biotin-11-ATP/UTP/CTP/GTP (PerkinElmer). After incubation, total RNA was extracted and fragmented by treatment with 0.2 M NaOH for 10 minutes on ice. Biotinylated nascent RNAs were then purified using streptavidin-coated M-280 beads (Invitrogen, #112.06D). Enzymatic treatments included the use of RppH (NEB, #M0356S) to remove the 5' cap, and PNK (NEB, #M0201) to repair the triphosphate and 5' hydroxyl ends. Adapter ligation and reverse transcription were performed using SuperScript III (Invitrogen, #56575), followed by PCR amplification. The final libraries were size-selected using AMPure XP beads (Beckman Coulter, #A63882) and sequenced on the Illumina NovaSeq 6000 platform with a single-read configuration.

**PRO-seq data analysis:**
Cutadapt (v1.14) and Trimmomatic (v0.32) were used to trim the raw FASTQ data (*4, 44*), and then Bowtie (v1.1.2) (*5*) was used to align the data on the human (hg38) or Drosophila (dm3) genome. The single nucleotide ends of aligned reads were made with BEDTools (v2.28) with genomecov (*11*) as bedgraph format. The number of reads mapped to the spike-in dm3 genome was used to normalize the bedgraph data before it was changed to bigwig data for further analysis. To determine the accurate TSS for each gene, we separated the mapped reads from PRO-seq by strand and individually called peaks using MACS2 (v1.2) (*9*) with the following parameters: --nomodel, --extsize 200 and –narrow to identify narrow peaks. We merged the common peaks from each sample for each stranded peak and used the BedTools command to annotate the peaks against mouse Gencode (v43). Peaks with a distance >100nt from an annotated TSS region were discarded to avoid multiple assignments to the same promoter. When multiple TSS from the same reference were detected in the same PRO-seq peak, we selected the most upstream isoform for analysis. Also, when a gene has multiple isoforms, the distance between these TSS was > 350nt and they were assigned to different PRO-seq peaks, we kept all isoforms for analysis. Next, we calculated the RPKM values in the promoter region (-50 nt from TSS to + 300 nt) and the region corresponding to the gene body (+301 nt from TSS to TES) for all transcripts used in the analysis (n = 13018). The read counts in the gene body region were used to calculate and determine the differential gene expression (DEG) between conditions using DESeq (*29*). Differentially expressed genes

were identified by using a 1.25-fold change cutoff (FDR<0.05) and applying a minimum gene expression filter for the final set of differentially expressed protein-coding genes (1 TPM in both replicates in at least one of the conditions). Gene set enrichment analysis (GSEA, v4.3.2) was performed on the Wald ranked list of genes using the weighted enrichment method.

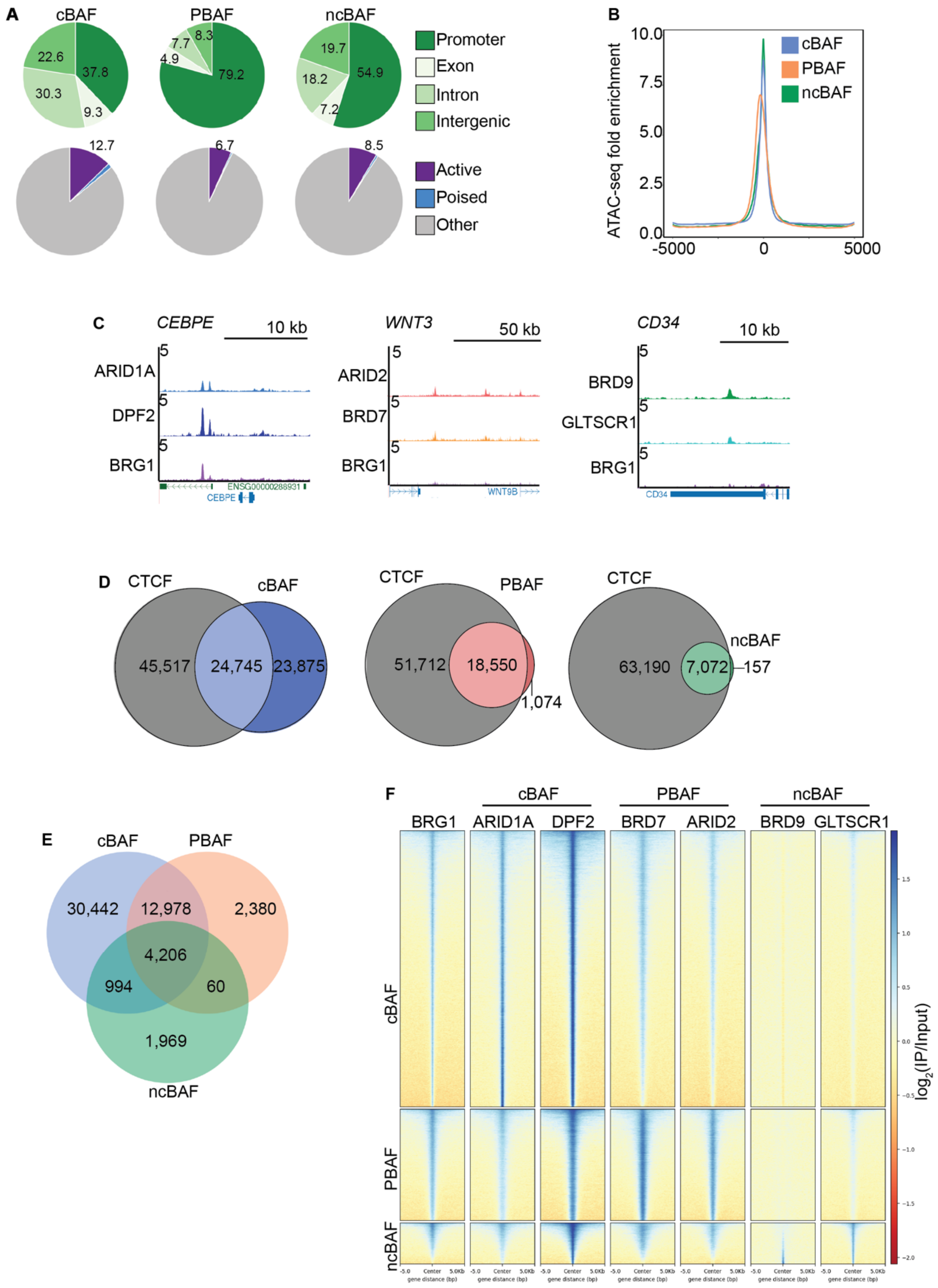


**Supplementary Figure 1. (A)** Pie charts of the genomic localization (*top*) and enhancer annotation (*bottom*) for the different BAF complexes. **(B)** Density plot of the ATAC-seq signal at the peaks bound by the different BAF complexes. The plot is peak-centered. **(C)** UCSC genome browser tracks at select loci that are bound by cBAF (*left*), PBAF (*middle*), or ncBAF (*right*). Representative tracks from one donor are shown, normalized to counts per million (CPM). **(D)** Area-proportional Venn diagrams of the overlap of CTCF ChIP-seq peaks with the peaks bound by the different BAF complexes. **(E)** Venn diagram of the overlap of the peaks of the cBAF, PBAF, and ncBAF complexes. **(F)** Heatmap of all peaks identified for the cBAF (ARID1A+ or DPF2+), PBAF (BRD7+ or ARID2+), and ncBAF (GLTSCR1+ or BRD9+) complexes**.**

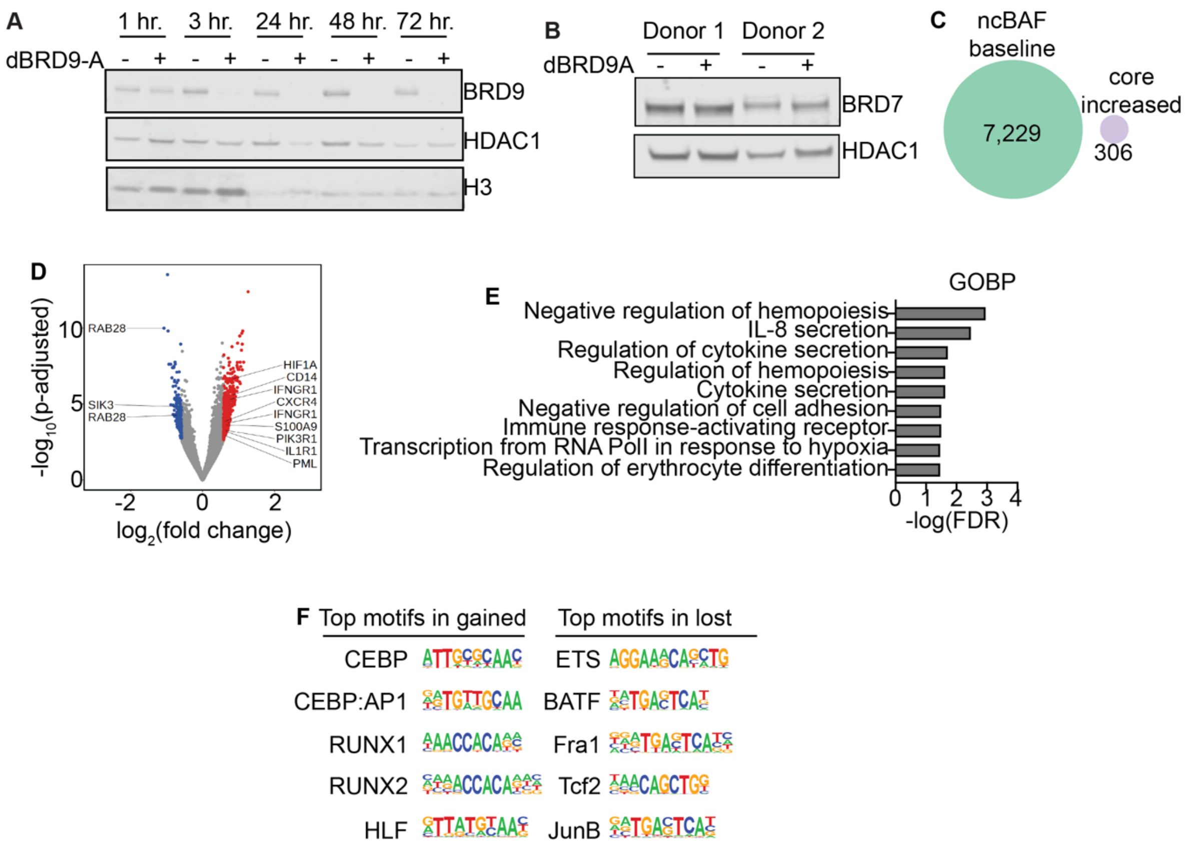


**Supplementary Figure 2. (A)** Time-course western blot of nuclear HUDEP-2 extracts after a single dose of dBRD9-A or vehicle control. H3 and HDAC1 loading controls are also shown. **(B)** Western blot of BRD7 in HSPC after 48 hours of dBRD9-A treatment. **(C)** Venn diagram of the overlap of ncBAF baseline peaks and peaks with a significant increase in core (BRG1) binding with loss of BRD9. **(D)** Volcano plot of the chromatin accessibility changes in HSPC treated for 72 hours with dBRD9-A or vehicle. Significant (p-adj < 0.05 and absolute($log_2$ (fold change)) > 1.5) differentially accessible regions are colored in red or blue. **(E)** Bar plot of select significant gene ontology biological processes that are enriched in the regions with significant increases in accessibility. **(F)** Topmost motifs for the transcription factors that are enriched in regions with gained (*left*) or decreased (*right*) accessibility with dBRD9-A treatment.

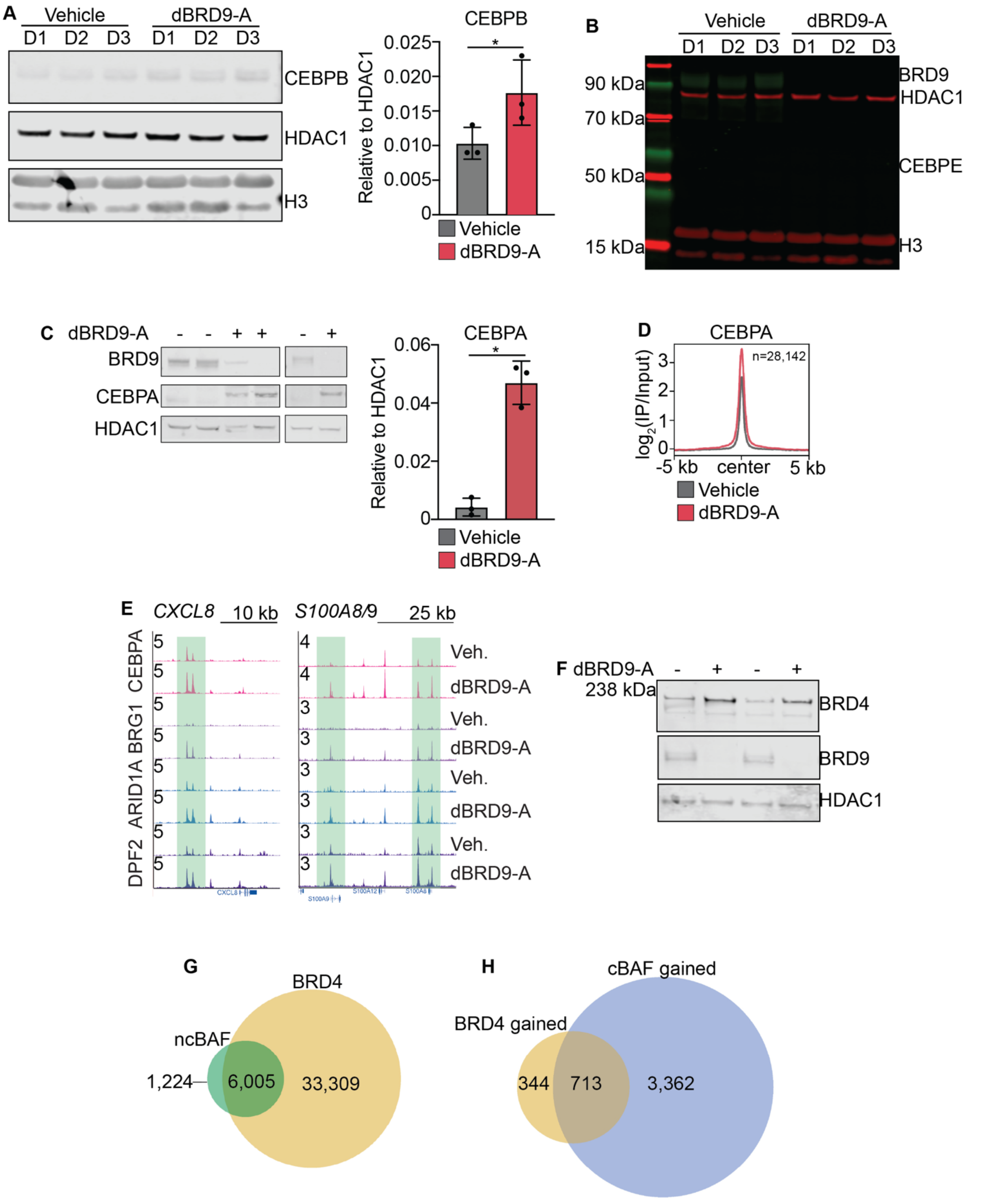


**Supplementary Figure 3. (A)** Western blot analysis and quantification of CEBPB in nuclear extracts from HSPC treated for 72 hours with dBRD9-A or vehicle control. HDAC1 and H3 loading controls are also shown (n=3 biological replicates). **(B)** Western blot of BRD9, CEBPE, HDAC1, and H3 in HSPC (n=3 biological replicates). **(C)** Western blots of BRD9, CEBPA, and HDAC1 in nuclear extracts from HSPC treated with dBRD9-A or vehicle control for 24 hours. Signal of CEBPA was normalized to HDAC1 loading control (*right*) (n=3 biological replicates). **(D)** Density plot of CEBPA signal at regions with a significant increase in CEBPA binding with loss of BRD9. **(E)** UCSC genome browser tracks of loci with significant increases in both CEBPA and cBAF binding with loss of BRD9. Tracks are from one representative donor and normalized to CPM. **(F)** Western of BRD4 and BRD9 in chromatin fraction from HUDEP-2 cells treated with dBRD9-A or vehicle. **(G)** Venn diagram of the overlap between the baseline ncBAF and BRD4 peaks. **(H)** Venn diagram of the overlap between genes with significant increases in cBAF and BRD4. Error bars represent mean with SD. Significance calculated using paired-t-test with Prism, p>0.05=ns, p ≤ 0.05=*, p≤0.01=**, p≤0.001=***, p≤0.0001=****.

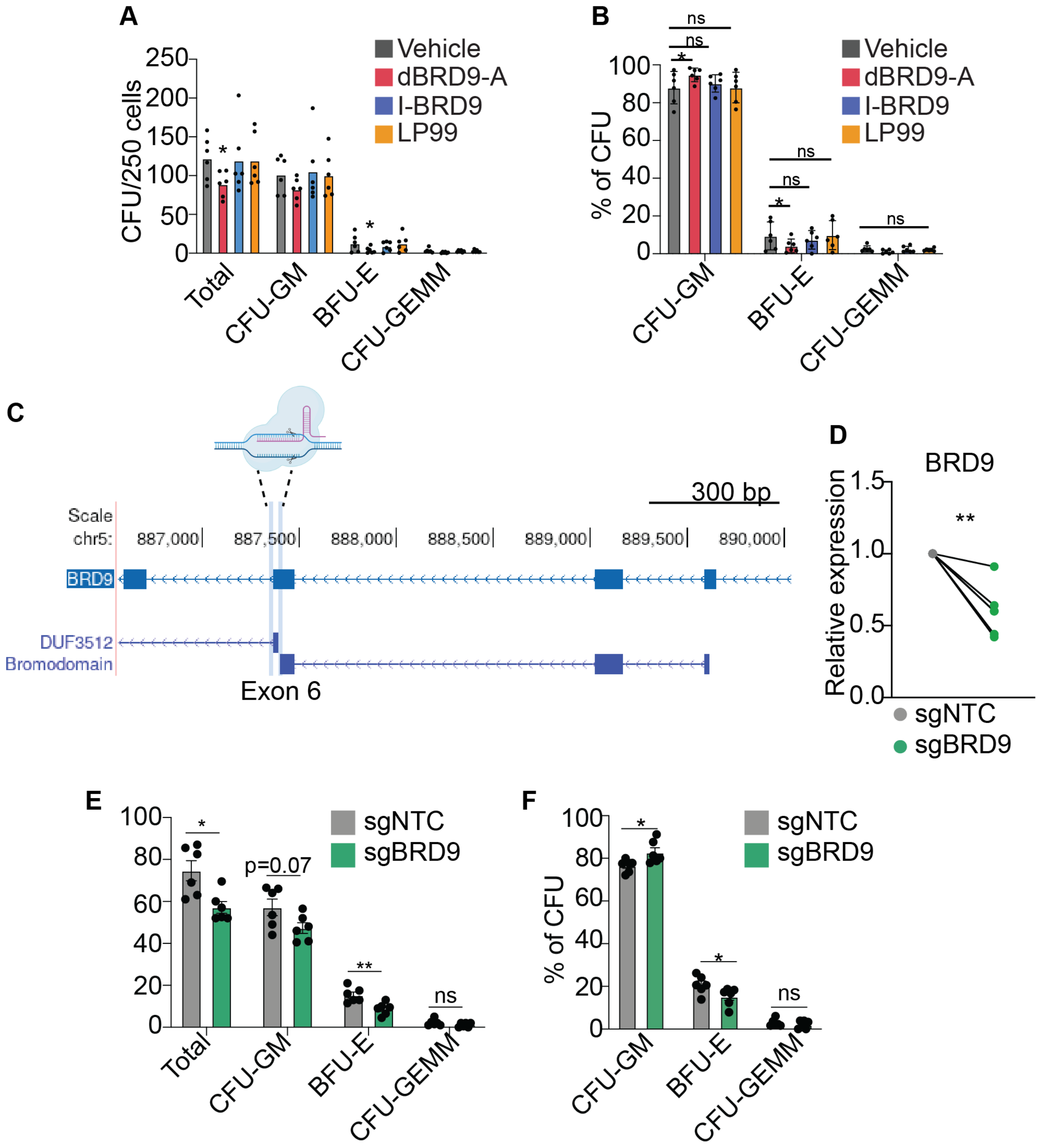


**Supplementary Figure 4. (A)** Bar plot of the number of total CFU enumerated from plating 250 HSPC treated with dBRD9-A, I-BRD9, or LP99 for 48 hours (n=6) and **(B)** the percentage of CFU types. **(C)** Schematic showing the location of the two sgRNA guides used to target exon 6 of BRD9. (**D**) Plot of the relative expression of BRD9 calculated by qRT-PCR in HSPC, 48 hours after CRISPR-Cas9 mediated KO of BRD9. (**E**) Bar plot of the number of total CFU enumerated from plating 200 sgNTC or sgBRD9 HSPC, 48 hours after electroporation (n=6) and **(F)** the percentage of CFU types. Error bars represent mean with SEM. Significance calculated using paired-t-test with GraphPad Prism, p>0.05=ns, p ≤ 0.05=*, p≤0.01=**, p≤0.001=***, p≤0.0001=****.

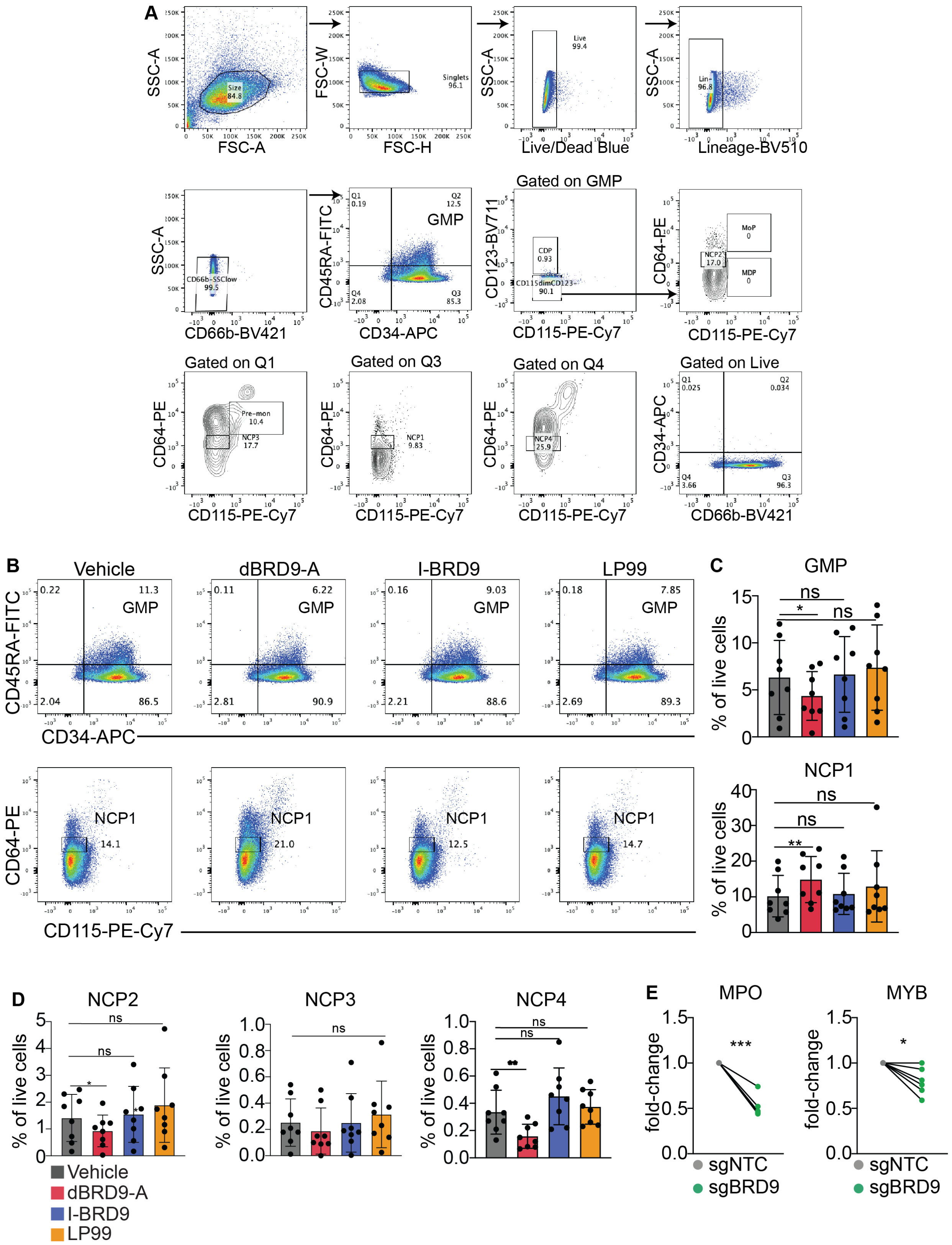
A
SSC-A
FSC-A
Size 84.8
FSC-W
FSC-H
Singlets 96.1
Live 99.4
Live/Dead Blue
Lin- 96.8
Lineage-BV510
CD66b-SSClow 99.5
CD66b-BV421
CD45RA-FITC
CD34-APC
Q1 0.19
Q2 12.5
GMP
Q4 2.08
Q3 85.3
Gated on GMP
CD123-BV711
CD115-PE-Cy7
CDP 0.93
CD115dimCD123- 90.1
CD64-PE
MoP 0
NCP2 17.0
MDP 0
Gated on Q1
Pre-mon 10.4
NCP3 17.7
Gated on Q3
NCP1 9.83
Gated on Q4
NCP4 25.9
Gated on Live
Q1 0.025
Q2 0.034
Q4 3.66
Q3 96.3
CD66b-BV421
B
Vehicle
dBRD9-A
I-BRD9
LP99
0.22
11.3
2.04
86.5
0.11
6.22
2.81
90.9
0.16
9.03
2.21
88.6
0.18
7.85
2.69
89.3
NCP1
14.1
21.0
12.5
14.7
C
GMP
% of live cells
ns
*
NCP1
**
D
NCP2
NCP3
NCP4
Vehicle
dBRD9-A
I-BRD9
LP99
E
MPO
MYB
fold-change
***
sgNTC
sgBRD9

**Supplementary Figure 5. (A)** Gating strategy used to identify neutrophil cell progenitors (NCP) and GMP. **(B)** Representative flow-cytometry plots of cell staining for GMP (*top*) and NCP1 (*bottom*) in HSPC treated with dBRD9-A, I-BRD9, LP99 or vehicle for 48 hours and cultured in pre-stimulation media and (**C**) summary bar plots of the frequencies as percentage of live cells. **(D)** Frequencies of NCP cells in HSPC treated with BRD9 chemical inhibitors or vehicle for 48 hours and measured with flow-cytometry (n=7). (**E**) Plots of the relative expression of MPO and MYB calculated by qRT-PCR in HSPC, 48 hours after CRISPR-Cas9 mediated KO of BRD9. Significance calculated using a paired t-test. Error bars represent mean with SD. Significance calculated using paired-t-test with Prism, $p>0.05$=ns, $p \leq 0.05$=*, $p\leq 0.01$=**, $p\leq 0.001$=***, $p\leq 0.0001$=****.

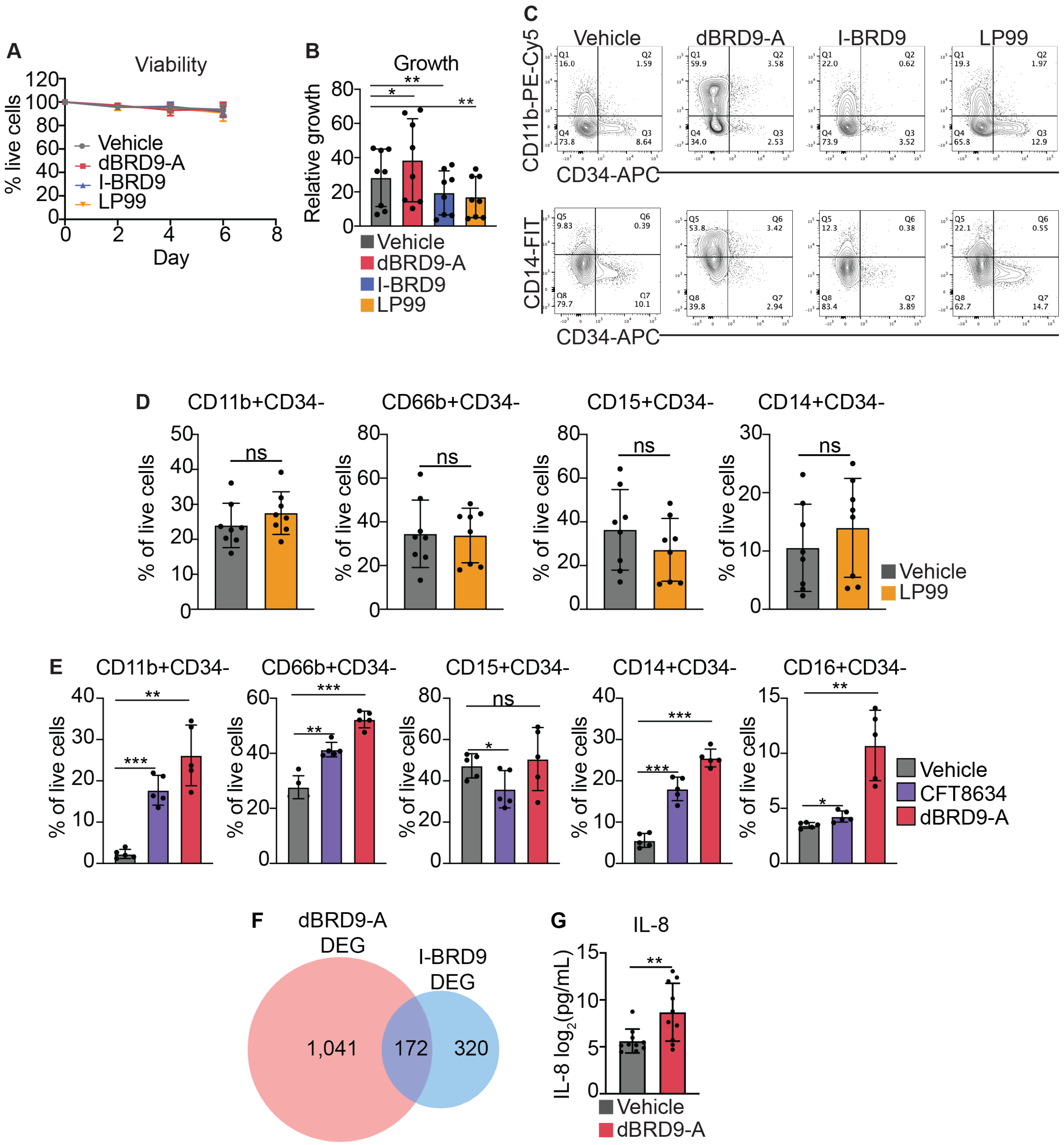


**Supplementary Figure 6. (A)** Graph of myeloid cell viability with treatment of dBRD9-A, I-BRD9, LP99, or vehicle control**. (B)** Graph of myeloid cell growth relative to day zero of culture. **(C)** Representative flow cytometry plots of CD34 vs. CD11b (*top*) and CD34 vs. CD14 (*bottom*) on day seven of myeloid differentiation. **(D)** Bar plots of the frequencies of different myeloid cell markers in myeloid cells treated for seven days with LP99 or **(E)** dBRD9-A, CFT8634, or vehicle control, as assessed by flow cytometry. **(F)** Venn diagram showing the overlap of genes that are differentially expressed with dBRD9-A or I-BRD9 treatment in myeloid cells compared to vehicle. **(G)** Graph of secreted IL-8 from conditioned media from dBRD9-A or vehicle-treated myeloid cells, as measured with a ProQuantum Immunoassay. Error bars represent mean with SD. Significance calculated using paired-t-test with Prism, $p>0.05$=ns, $p \leq 0.05$=*, $p \leq 0.01$=**, $p \leq 0.001$=***, $p \leq 0.0001$=****.

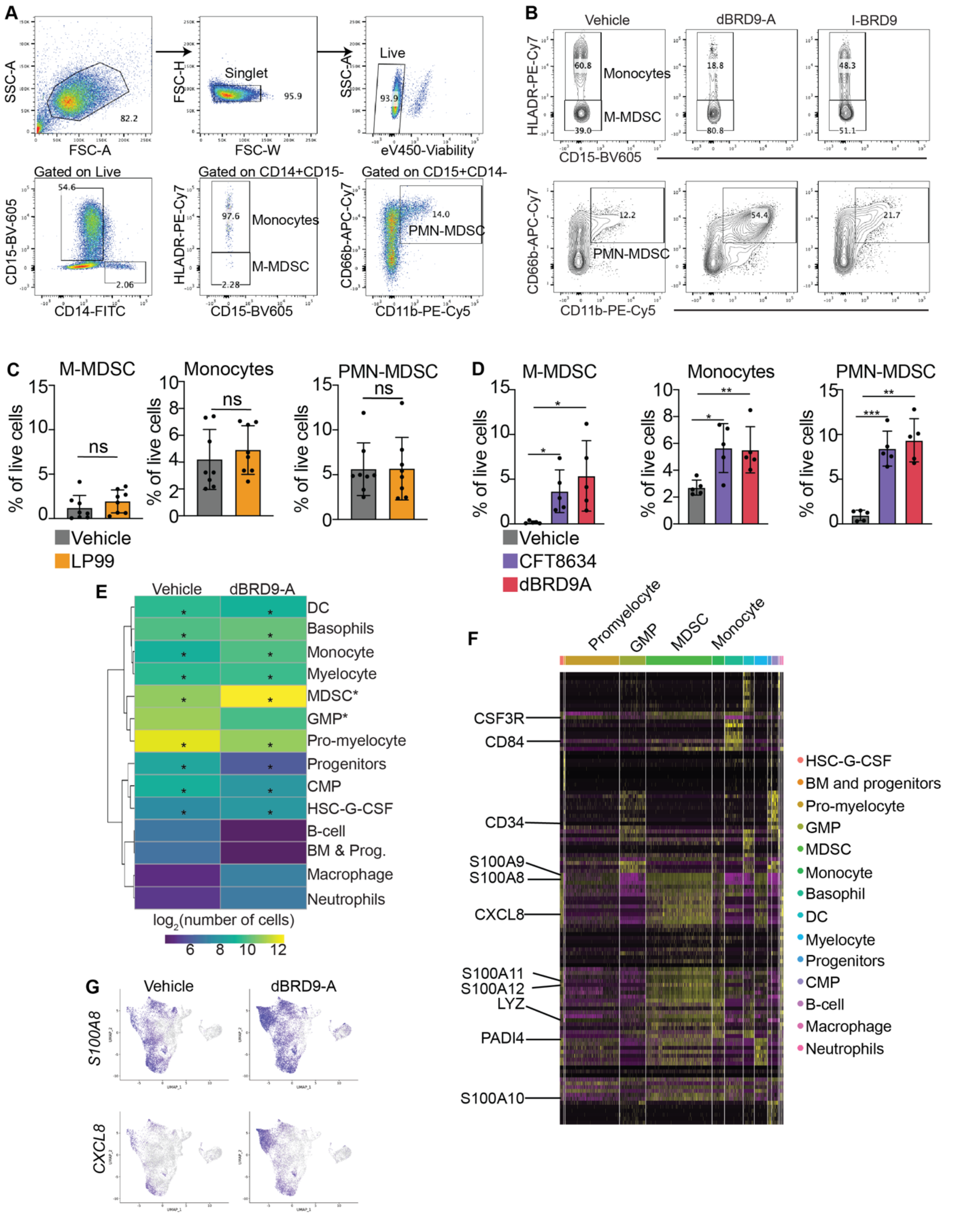


**Supplementary Figure 7. (A)** Gating strategy used to identify human M-MDSC and PMN-MDSC. **(B)** Representative flow-cytometry plots of monocytes, M-MDSC, and PMN-MDSC on day seven of myeloid differentiation. M-MDSC are gated on CD14+CD15- cells, and PMN-MDSC are gated on CD15+CD14- cells. **(C)** Barplot of the percentages of MDSC and monocytes as a frequency of live cells, from myeloid cells treated with LP99 or vehicle (n=8) or **(D)** vehicle, dBRD9-A, or CFT8634 (n=5). **(E)** Heatmap showing the number of each cell type within the vehicle or dBRD9-A myeloid cell population. Asterisks denote populations that are significantly enriched (FDR < 0.05) with loss of BRD9. **(F)** Heatmap showing the gene expression for each cell for the top cell identity genes for each cell type, annotated, including both dBRD9-A- and vehicle-treated cells. Select genes are labeled. **(G)** UMAP plots of select genes that are significantly enriched in MDSC compared to other cell types assessed. Error bars represent mean with SD. Significance calculated using paired-t-test with Prism, p>0.05=ns, p ≤ 0.05=*, p≤0.01=**, p≤0.001=***, p≤0.0001=****.

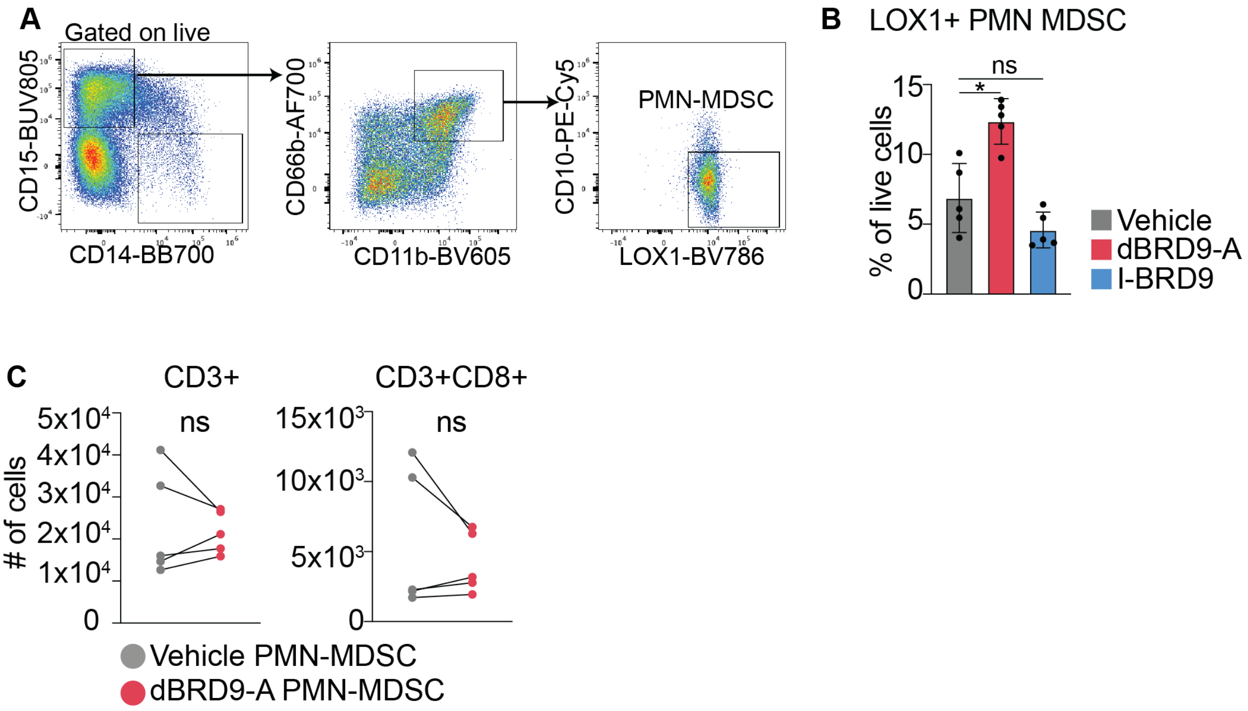


**Supplementary Figure 8: (A)** Gating scheme to identify LOX1+CD10- PMN-MDSC. **(B)** Bar plot of the percentage of LOX1+ PMN-MDSC in myeloid cells treated with vehicle, dBRD9-A, or I-BRD9. **(C)** Plots of the number of CD3+ T-cells (*left*) or CD3+CD8+ T-cells (*right*) after co-culture with PMN-MDSC that had been treated with dBRD9-A or vehicle (n=5). Error bars represent mean with SD. Significance calculated using paired-t-test with Prism, $p>0.05$=ns, $p \leq 0.05$=*, $p\leq0.01$=**, $p\leq0.001$=***, $p\leq0.0001$=****.

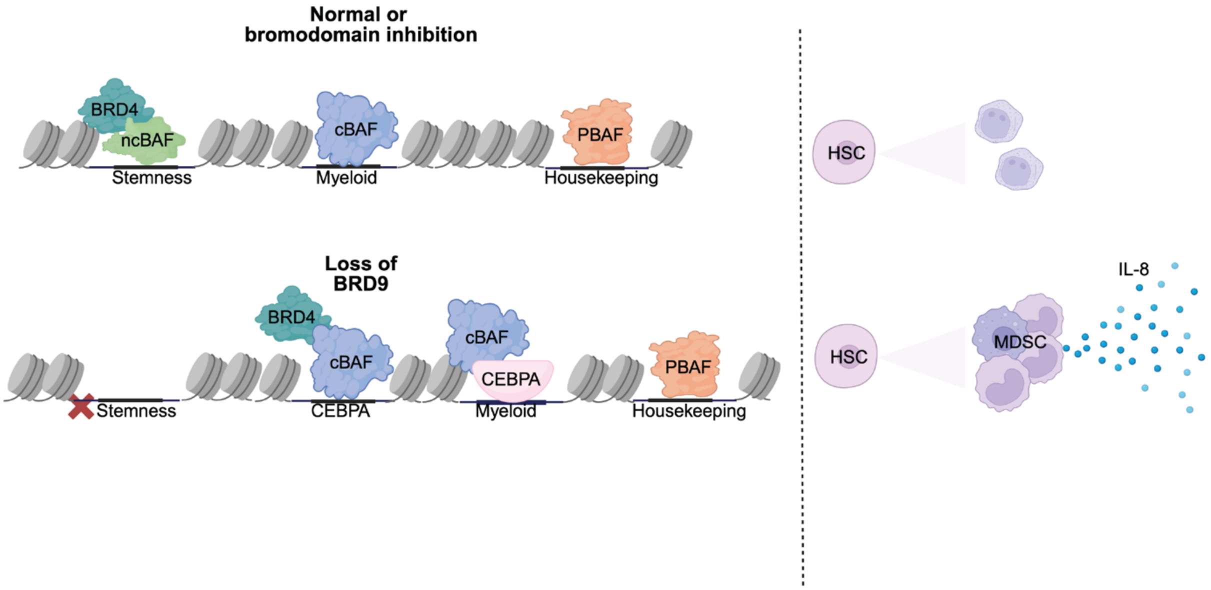


**Supplementary Figure 9. (A)** Graphical abstract depicting the role of BRD9 in regulating the expression of human granulocytic genes and the differentiation of suppressive myeloid cells.
